# Pretrauma cognitive traits predict trauma-induced fear generalization and associated prefrontal functioning in a longitudinal model of posttraumatic stress disorder

**DOI:** 10.1101/2024.03.11.584500

**Authors:** László Szente, Manó Aliczki, Gyula Y. Balla, Róbert D. Maróthy, Zoltán K. Varga, Bendegúz Á. Varga, Zsolt Borhegyi, László Biró, Kornél Demeter, Christina Miskolczi, Zoltán Balogh, Huba Szebik, Anett Szilvásy-Szabó, Anita Kurilla, Máté Tóth, Éva Mikics

**Author notes:** Equal contributions (Máté Tóth, Éva Mikics).

## Abstract

Posttraumatic stress disorder (PTSD) is a chronic psychiatric condition that develops in susceptible individuals exposed to traumatic stress, challenging clinicians to identify risk factors and mechanisms for mitigating vulnerability. Here we investigated behavioral predictors of high fear generalization, a core PTSD symptom, and its neural correlates longitudinally in rats. In a comprehensive behavioral test battery of emotional and cognitive function, pretrauma lower operant learning performance emerged as high predictor of fear generalization following trauma. Posttrauma operant training facilitated fear extinction, suggesting an overlap in neural circuits governing operant learning and fear expression. Neuronal activity mapping revealed significant changes in the medial prefrontal cortex (mPFC) in high fear generalizers, with alterations in CRH/VIP+ interneuron functioning. Silencing prefrontal *Crh* expression after fear memory consolidation enhanced mPFC activation and reduced fear expression, favoring resilience. These findings highlight operant learning and mPFC alterations as vulnerability markers and mediators of excessive fear generalization, with implications for prevention and targeted therapy in PTSD.

## Introduction

Posttraumatic stress disorder (PTSD) is a chronic psychiatric condition that emerges following overwhelming stressors, affecting a subset of individuals exposed to trauma with a prevalence ranging between 10-20% (Kessler et al., 1995; Breslau et al., 1998). Pretrauma risk factors and mechanisms determining individual vulnerability need to be identified for better preventions (in high-risk populations) and therapies targeting vulnerability mechanisms (Yehuda and LeDoux, 2007). Given the inherent challenges in measuring and testing premorbid conditions and their causal role in vulnerability in humans, longitudinal animal studies become essential to complement PTSD research conducted on high-risk human populations (Cohen et al., 2004; Richter-Levin et al., 2019).

While multiple symptoms emerge in PTSD (including hyperarousal, changes in mood and cognition, avoidance, intrusive memories), maladaptive core symptoms can be defined as intrusive/excessive fear memories, which are generalized to safe contexts and resistant to extinction (Rauch et al., 2006; Shin and Handwerger, 2009; Liberzon and Abelson, 2016). These characteristics also represent current clinically defined targets of therapies (i.e. enhanced fear extinction by exposure or other therapeutic interventions), which are promising but require further improvements (Singewald et al., 2015). Moreover, DSM-5 based diagnostic categories describe complex conditions with multiple symptoms (American Psychiatric Association, 2013), which needs deconstruction into biologically relevant neurobehavioral domains to clarify underlying mechanisms as it was proposed in the Research Doman Criteria (RDoC) framework (Insel et al., 2010). RDoC-based animal models provide the feasibility for this approach, and hence, they can imply translationally relevant targets (Anderzhanova et al., 2017; Richter-Levin et al., 2019). Therefore, we focused on these core symptoms as outcome measures with an additional advantage of their high evolutionary conservation across species, hence, maximizing translational validity of our findings (Flandreau and Toth, 2018; Singewald and Holmes, 2019).

We aimed to characterize individual predisposing affective and cognitive traits pretrauma, which reliably predict trauma-induced persisting fear generalization, and characterized underlying neural mechanisms using an unbiased top-down approach, i.e. from phenotype to neural circuits, and to cellular-molecular levels. We used a modified Pavlovian fear conditioning paradigm with long-term outcomes and contrasted vulnerable and resilient subpopulations (i.e. high vs. low fear generalizers) to detect vulnerability markers and related mechanisms.

We put significant emphasis and performed detailed characterization on predisposing cognitive traits since there is limited evidence on how specific cognitive traits/subdomains contribute to PTSD vulnerability. Lower cognitive performance has previously been reported in PTSD (DiGangi et al., 2013; Schultebraucks et al., 2021), however, its presence pretrauma as causal mediator is not clarified. Similarly, structural and functional alterations of the hippocampus and prefrontal regions as potential mediators of altered cognition are documented in PTSD, however, their pretrauma causal role is not clarified (Kasai et al., 2008; Pitman et al., 2012). Moreover, the detailed understanding of cellular and network level alterations of these regions in the development of PTSD, particularly related to specific neurobehavioral symptoms (e.g. fear generalization), is needed for better prevention and treatment options (Lopresto et al., 2016; Asok et al., 2018).

In summary, we identified the operant cognitive domain as the highest predictor for trauma-induced fear generalization with associated brain activity and gene expression changes. Medial prefrontal cortical (mPFC) network regulated by CRH/VIP+ interneurons emerged as major determinant of trauma-related excessive fear generalization, which is detectable pretrauma by lower operant learning performances. Furthermore, enhanced recruitment of this circuitry by additional operant training can enhance the extinction of generalized fear potentially via plasticity-related changes.

## Results

### Unpredictable footshock induces lasting fear memory with persisting fear generalization in vulnerable subpopulation

We first aimed to establish and characterize an experimental paradigm that allows for identification of vulnerable and resilient subpopulations following trauma. To achieve this, we applied a single series of electric footshocks to rats as traumatic stimulus and assessed their fear responses a month later in the same, trauma-associated context as well as in an altered/safe context, that allowed for identification of vulnerable individuals based on their core PTSD symptoms i.e. enhanced fear generalization of contextual fear (Fig.1a and b).

**Fig. 1.**
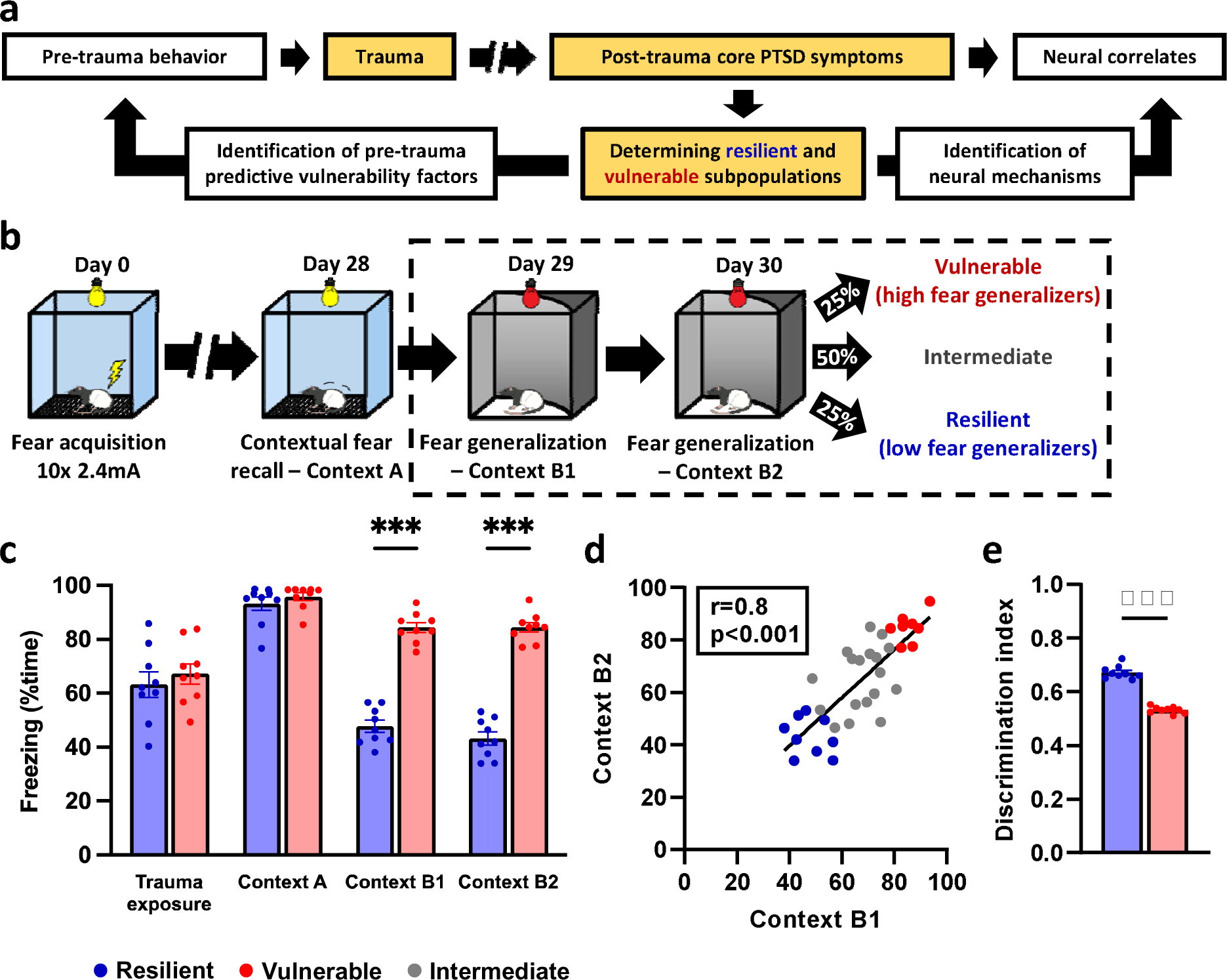
Identification of resilient and vulnerable subpopulations. **a** Experimental design. Data are shown from experimental stages highlighted in yellow. **b** Animals were exposed to electric footshocks on Day 0, followed by a brief exposure to the same context (Context A) on Day 28, and two exposures to an altered/safe context (Context B) on the next two consecutive days (Day 29 and 30). Based on their average freezing levels in context B1 and B2 tests, subjects were sorted into resilient and vulnerable groups as lower and upper quartiles (25-25%), respectively. Subjects in the intermediate two quartiles were defined as an ‘intermediate’ group and were not analyzed, except when correlation analyses were performed on the whole population. **c** Resilient and vulnerable groups displayed marked difference in their freezing response in Context B1 and B2, although their freezing responses were similar during fear acquisition (i.e. trauma exposure) and during exposure to the traumatic/conditioned context (Context A). **d** Individual freezing levels exhibited in Context B1 and B2 were highly consistent across the population indicated by a significant positive correlation. **e** Similarly to freezing levels in Context B, context discrimination index (freezing in Context B / freezing in Context A) as an indicator of fear generalization was also markedly different between resilient and vulnerable groups. All data are shown as mean ± S.E.M. ***p<0.001; Unpaired t-test, Mann-Whitney U test, and Pearson correlation.

Unpredictable electric footshocks induced freezing response in all subjects that gradually increased across the conditioning session with the number of footshocks (F_shock_(1,41)=245.340, p<0.001; F_time_(2,82)=10143.92, p<0.001; Fig.1c). This experience resulted in lasting fear memory indicated by marked freezing responses during re-exposure to the trauma-associated context (Context A: CtxA) 4 weeks later (U=0, p<0.001, compared to non-shocked controls; Fig.1c). This contextual fear recall showed minimal variability in the population indicating strong memory recall of the trauma in all subjects (Fig.1c). In contrast, exposure to an altered/neutral ‘safe’ context (Context B: CtxB) on two consecutive days precipitated a highly variable freezing response across subjects (Fig.1c and d), which was stable within subjects across test days as freezing levels on consecutive testing days in CtxB showed strong positive correlation (r=0.797, p<0.001; Fig.1d) indicating that trauma-induced fear generalization is an individual trait-like characteristic. Our meta-analysis of fear responses of all experimental cohorts presented in our study confirmed the above findings: strong memory encoding with highly variable generalization levels (Supplementary Fig.1) that also showed bimodal distribution tendencies: unimodal distribution was rejected (Excess mass=0.077, p<0.001), however, the presence of more than 2 modes could not be shown (Excess mass=0.028, p=0.3).

Latter indicated diverging outcome trajectories, hence, supporting the validity to discriminate vulnerable and resilient subpopulations posttrauma.

Based on the marked variability and individual trait-like characteristic of generalized fear, this model allowed the separation of vulnerable and resilient subpopulations exhibiting stable high and low fear generalization, respectively (upper and lower quartiles based on freezing response in CtxB; CtxB1: F_vuln_(1,16)=157.4, p<0.001; CtxB2: F_vuln_(1,16)=188.4, p<0.001; Fig.1c; contextual fear discrimination index (CtxB/CtxA): U=0, p<0.001; Fig.1e). Importantly, freezing response during footshock exposure did not differ between vulnerable and resilient subpopulations (F_vuln_(1,16)=0.422, p=0.525; F_vuln*time_(2,32)=0.878, p=0.425; Fig.1c), suggesting that acute sensitivity/reactivity to aversive stimuli is not a significant determinant of fear generalization. The subpopulations also showed similar freezing responses during contextual fear recall in CtxA (U=30, p=0.386; Fig.1c), indicating strong memory encoding of traumatic experience in both groups. Accordingly, differences in fear generalization of vulnerable and resilient subpopulations did not originate from learning or memory recall differences. These findings were also confirmed by our meta-analysis containing all cohorts of the present study showing similar freezing response of vulnerable and resilient subpopulations during footshock and CtxA exposures (Supplementary Fig.1a-c). Although fear generalization was the most prominent difference between vulnerable and resilient subpopulations, slower extinction curve of generalized fear in vulnerable subjects was also apparent in our meta-analysis (F_vuln*time_(7,1176)=3.491, p=0.001; Supplementary Fig.1d). These findings above indicate that our experimental paradigm is appropriate for identifying and examining vulnerable subpopulations that exhibit key symptoms of PTSD after experiencing trauma, i.e. fear generalization and extinction deficit. Consequently, as detailed later, our model enabled us to conduct longitudinal investigations and identify *pretrauma* emotional and cognitive characteristics that could specifically forecast posttrauma fear generalization and difficulties in fear memory extinction.

### Pretrauma affective traits are weak predictors of fear generalization

First, we assessed pretrauma affective/anxiety-like traits as PTSD has been primarily considered as anxiety disorder, but recently it has also been excluded from this nosology group (DSM-IV vs DSM-5; (American Psychiatric Association, 1994, 2013). In our study, retrospective analysis of pretrauma anxiety traits showed that there were no discernible group differences in the two less aversive tests: neither the behavior observed on the elevated plus maze (%time in open arm: F_vuln_(1,16)=0.145, p=0.708; z-score: F_vuln_(1,16)=0.015, p=0.900; Fig.2c) nor in the open field (%time in center: F_vuln_(1,16)=0.275, p=0.606; z-score: F_vuln_(1,16)=0.055, p=0.816; Fig.2d) showed differences between vulnerable and resilient groups. In contrast, the vulnerable group exhibited higher avoidance in the two highly aversive tests, i.e. in the light-dark box (%time in light compartment: F_vuln_(1,16)=3.784, p=0.070; z-score: F_vuln_(1,16)=8.410, p=0.010; Fig.2b), and during predator odor exposure test, latter on a trend level (freezing: F_vuln_(1,16)=2.615, p=0.125; F_vuln_(1,16)=3.846, p=0.067; Fig.2e). Pretrauma acoustic startle reactivity was not different between resilient and vulnerable groups (F_group_(1,16)=0.720, p=0.408; F_intensity*group_ (4,64)=2.124, p=0.088; Fig.2f). These results indicate that anxiety-like behavior under higher threats has significant, but low predictive power for fear generalization following traumatic stress exposure.

**Fig. 2.**
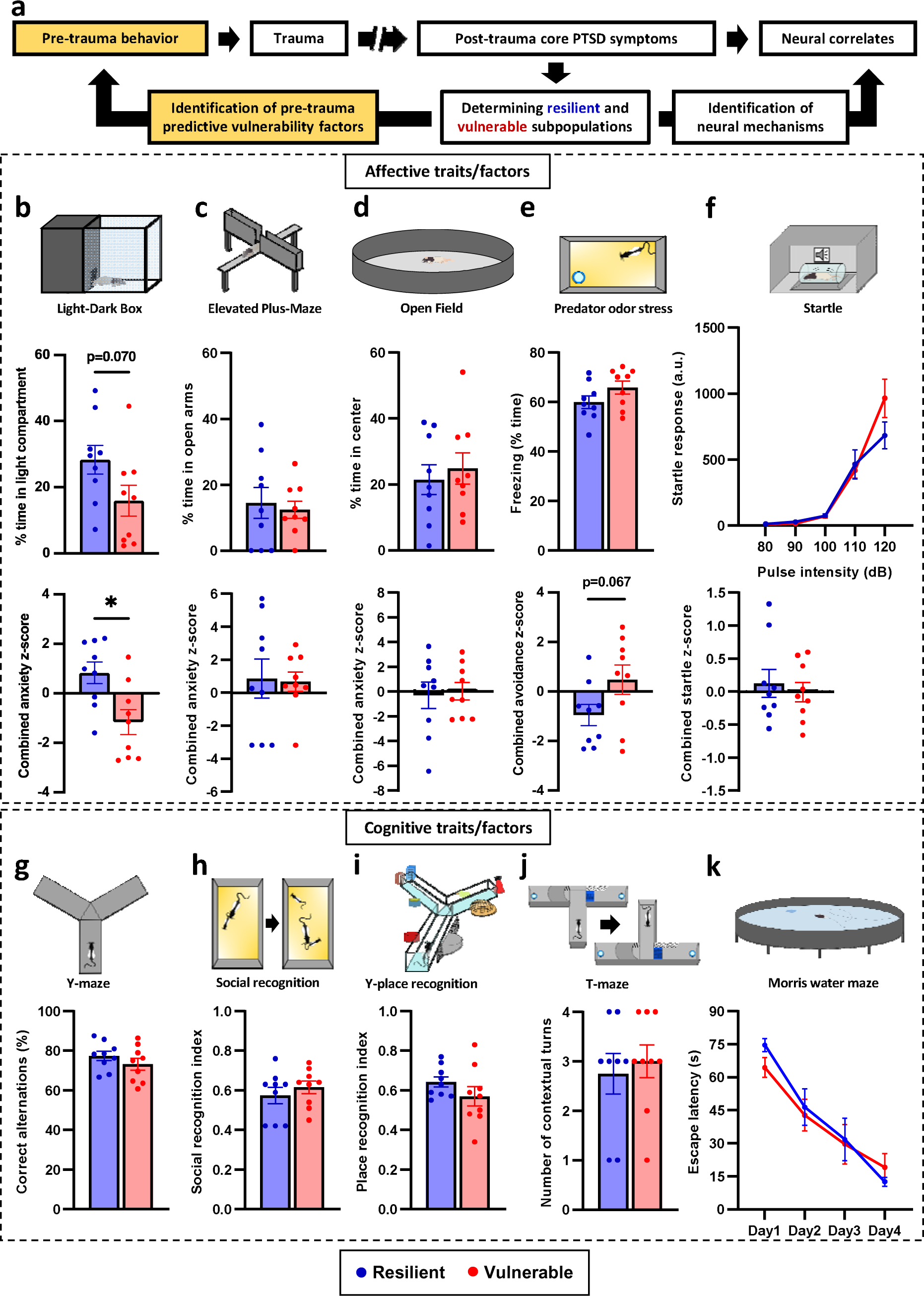
Identification of pre-trauma vulnerability factors by assessment of affective (upper box: b-f) and cognitive (lower box:g-k) traits. **a** Experimental design. Data are shown from experimental stages highlighted in yellow. **b** Anxiety measures in the Light-Dark Box showed pretrauma difference between groups as shown by major parameter (time% in aversive light compartment; middle panel) and combined z-score (lower panel). **c-d** Anxiety measures in the Elevated Plus Maze and Open Field showed no group differences. **e** The Predator odor test showed a trend for group difference indicated by the combined z-score (lower panel). **f** Startle reactivity was not different between groups. **g** Alternations in the Y-maze as an index of working memory did not reveal group difference. **h** Social recognition and **i** place recognition (as indexes of social and contextual memory) did not differ between groups. **j** Contextual vs. habitual strategies in the T-maze were not different between groups. **k** Spatial learning in the Morris Water Maze was also similar between groups. All data are shown as mean ± S.E.M. *p<0.05, **p<0.01; One-way ANOVA, repeated measures ANOVA, or Mann-Whitney test.

### Pretrauma cognitive vulnerability factors I.: social and contextual learning, habitual- contextual strategy, and working memory traits are not predictive for fear generalization

Lower cognitive performance has been documented in PTSD, however its contribution to PTSD vulnerability and related neurobiological alterations remain to be elucidated. Therefore, we performed a series of pretrauma cognitive tests to assess their predictive power for fear generalization outcomes after trauma.

Performance in several cognitive tasks revealed no prediction for subsequent fear generalization. Accordingly, the number of correct/spontaneous alternations, a measure of short-term spatial working memory in the Y-maze test did not differ between the resilient and vulnerable groups (F_vuln_(1,16)=1.182, p=0.293; Fig.2g). Similarly, social recognition (U=28.00, p=0.289; Fig.2h) and place recognition indices (F_vuln_(1,16)=1.763, p=0.202; Fig.2i), representing intermediate/short-term memory in the social and Y-place recognition tests, was not different between groups. Spatial learning performance in the Morris water maze (F_days_(3,42)=35.612, p<0.001; F_vuln_(1,14)=0.104, p=0.751; F_vuln*days_(3,42)=0.816, p=0.491; Fig.2k), and habitual vs. contextual strategies in the T-maze tests (U=32, p=0.73; Fig.2j) were also similar between groups. These findings indicate that memory-related (working memory, encoding, recall) function and ability to use contextual cues are not predictive for trauma-related fear generalization.

### Pretrauma vulnerability factors II.: operant learning and cognitive flexibility traits are predictive for fear generalization

Vulnerable subjects required significantly more days to learn an operant task based on a visual cue (U=13, p=0.040; Fig.3a-b). This link between pretrauma operant performance and vulnerability was also confirmed on a populational level (including intermediate quartiles) by a positive correlation between the number of days to learn task criteria and fear generalization (r=0.462, p=0.003; Fig.3c). In a separate cohort, we found the same effect using a more complex operant learning task (Fig.3f): vulnerable group required more days to learn the task (F_vuln_(1,14)=5.923, p=0.028; Fig.3g), and showed less accuracy during the task (F_days_(4,56)=5.362, p<0.001; F_vuln_(1,14)=5.620, p=0.032; F_vuln*days_(4,56)=3.739, p=0.009; Fig.3i).

**Fig. 3.**
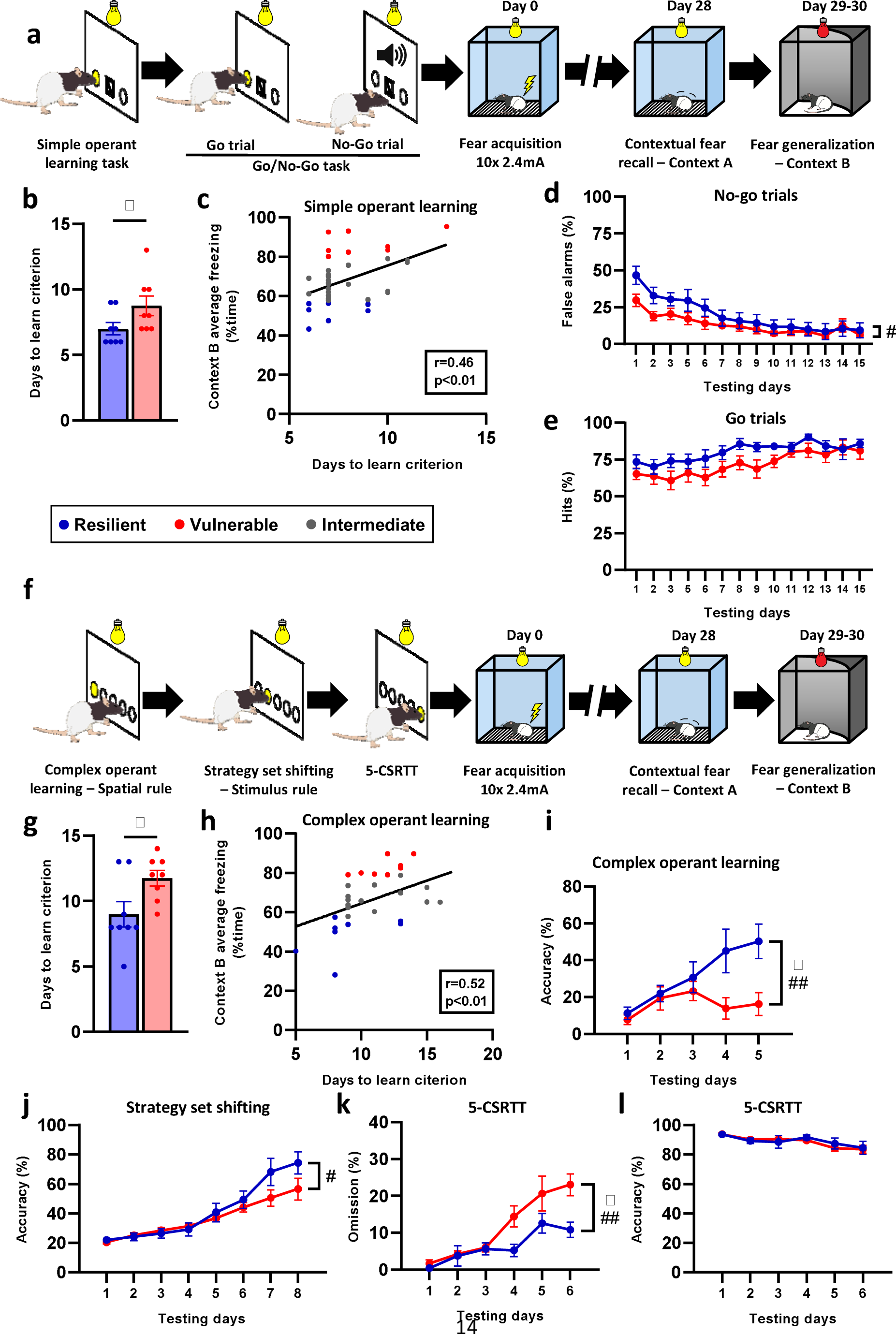
Pre-trauma operant learning ability predicts vulnerability/high fear generalization. **a** Experimental design of pre-trauma screening of simple operant learning and Go/No-Go tasks. **b** Simple operant task was significantly predictive as vulnerable subjects needed more days to learn the task criterion (>80% accuracy in 2 consecutive days) compared to resilient subjects. **c** ‘Days to learn’ criterion also significantly correlated with freezing level in Context B. **d** Go/No- Go task did not show significant group difference, although false alarms (NoGo errors) were somewhat lower at the early phase of the task (significant group*day interaction). **e** Go responses were similar between groups. **f** Experimental design of another pre-trauma screening of a more complex operant learning task extended with a Strategy Set Shifting Task and 5-Choice Serial Reaction Time Task (5-CSRTT). **g** Similarly to the simple operant learning task, vulnerable subjects needed significantly more days to learn the criterion of the more complex operant task compared to the resilient subjects. **h** Likewise, ‘Days to learn’ criterion correlated with freezing level in the Context B. **i** In addition to criterion completion difference, response accuracy showed a persistently lower performance in the vulnerable group. **j** The strategy set-shifting task also indicated lower performance in the vulnerable group indicated by slower increase in accuracy (group*day interaction). **k** Omissions, but not **l** accuracy in the 5-choice serial reaction time test was higher in the vulnerable group (group*day interaction). All data are shown as mean ± S.E.M. *p<0.05, **p<0.01, ***p<0.001, significant difference from the resilient group; #p<0.05, ##p<0.01, significant interaction; Unpaired t-test, Mann-Whitney U test, One-way ANOVA with Tukey’s post hoc test, or Repeated measures ANOVA.

Similar to the previous cohort, days required to learn the task correlated positively with fear generalization on a populational level (r=0.522, p=0.003) (Fig.3h).

Inhibitory functions assessed in the subsequent Go/No-Go task (in the first cohort, following operant task) were not predictive for fear generalization. Although false alarms (i.e. No-Go errors) showed a significant day*group interaction (F_days_(13,182)=23.13, p<0.001; F_vuln_(1,14)=1.586, p=0.228; F_vuln*days_(13,182)=2.070, p=0.017; Fig.3d), a trend-like group difference in Go responses (F_days_(13,182)=5.599, p=0.001; F_vuln_(1,14)=4.257, p=0.058; F_vuln*days_(13,182)=0.649, p=0.810; Fig.3e) indicated that the vulnerable group responded less frequently for both type of stimuli, and not specifically inhibitory responses were affected. Latter implies a more inhibited response tendency (when a novel acoustic stimulus was presented), rather than a difference in inhibitory control.

In the case of the strategy set-shifting task, vulnerable subjects shifted their response and reached accuracy slower compared to resilient subjects (F_days_(7,98)=40.95, p<0.001; F_vuln_(1,14)=1.148, p=0.302; F_vuln*days_(7,98)=2.583, p=0.017; Fig.3j) implying less cognitive flexibility. In the subsequent 5-choice serial reaction time task (5-CSRTT, measuring attentive function), vulnerable subjects showed lower performance reflected in the rate of omissions (F_days_(5,65)=21.81, p<0.001; F_vuln_(1,13)=5.487, p=0.040; F_vuln*days_(5,65)=3.352, p=0.009; Fig.3k), but not in accuracy (F_days_(5,65)=3.404, p=0.008; F_vuln_(1,13)=0.521, p=0.483; F_vuln*days_(5,65)=1.015, p=0.416; Fig.3l) suggesting somewhat lower attentional capacity, although it is inconclusive whether this is driven by inhibited response tendency (also present in the Go/No-Go task). In summary, pretrauma slower operant learning and cognitive inflexibility in strategy set-shifting showed the strongest association with high levels of fear generalization posttrauma.

### Posttrauma cognitive training enhances the extinction of generalized fear in association with plasticity-related gene expression changes in prefrontal and hippocampal regions

Based on the strong prediction of pretrauma operant learning traits for trauma-induced fear generalization, we hypothesized a potential cross-talk between cognitive performance and fear generalization, which is potentially mediated by overlapping neural circuits with functional differences between vulnerable and resilient groups. Therefore, we tested whether enhanced recruitment of these overlapping circuits, by means of posttrauma operant training as a potential therapeutic intervention, have significant impact on fear generalization and extinction. To address this, rats were tested for fear generalization after trauma and then underwent operant training procedures, after which they were re-tested for generalization and extinction.

We sorted resilient and vulnerable groups into ‘trained’ and ‘yoked control’ subgroups with counterbalanced freezing levels in CtxB (F_vuln_(1,30)=69.19, p<0.001; F_training_(1,30)=0.131, p=0.720; Fig.4a-b). Freezing response during trauma exposure (F_vuln_(1,30)=3.772, p=0.061; F_training_(1,30)=0.837, p=0.367; F_vuln*training_(1,30)=0.116, p=0.734; Supplementary Fig.2a) and CtxA exposure (F_vuln_(1,30)=0.322, p=0.574; F_training_(1,30)=0.160, p=0.692; F_vuln*training_(1,30)=3.036, p=0.073; Supplementary Fig.2b) did not differ significantly between groups prior to cognitive training. Fear memory was still robust after the operant learning task indicated by freezing response during post-training CtxA exposure, which was slightly lowered in the resilient following the operant training (F_vuln_(1,30)=12.10, p=0.001; F_training_(1,30)=0.066, p=0.798; F_vuln*training_(1,30)=0.003, p=0.956; Supplementary Fig.2c). In contrast to contextual recall, operant training accelerated the extinction curve in the vulnerable group (F_time_(3,45)=9.753, p<0.001; F_training_(1,15)=4.644, p=0.047; F_time*training_(3,45)=1.763, p=0.167; Fig.4d, right), which was not apparent in the resilient group, potentially due to a floor effect, i.e. resilient group exhibiting rather stable-low freezing levels (F_time_(3,45)=0.353, p=0.786; F_training_(1,15)=0.032, p=0.859; F_time*training_(3,45)=0.758, p=0.523; Fig.4d, left).

**Fig. 4.**
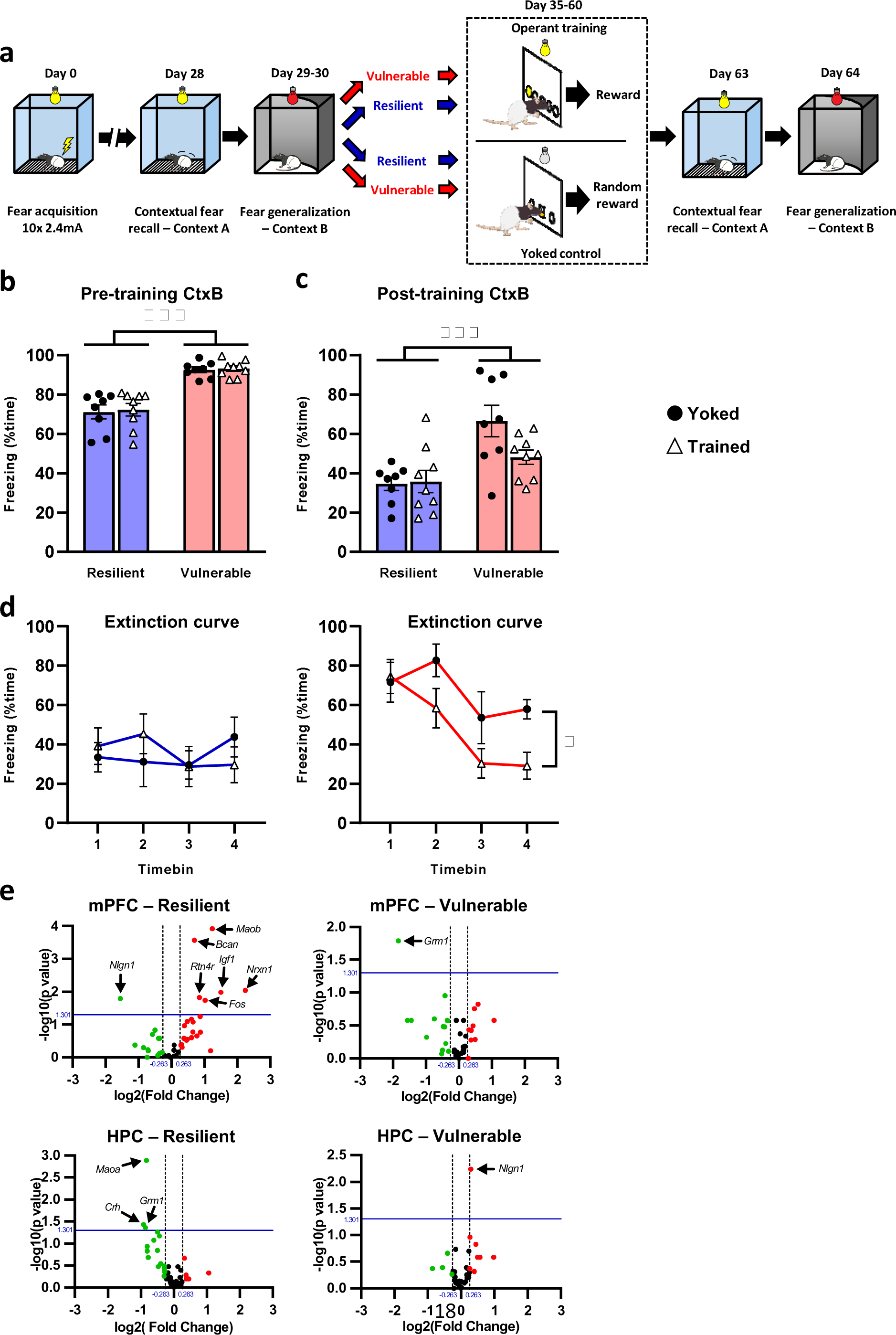
Post-trauma operant training facilitates extinction of fear generalization in vulnerable subjects. **a** Experimental design of post-trauma cognitive training and its impact on fear recalls. Subjects were sorted into resilient and vulnerable groups based on their freezing in Context B, then groups were further divided into trained and yoked (control) subgroups. Following a post- trauma operant cognitive training, fear responses were re-tested in context A and B. **b** Pre- training fear response in Context B was significantly higher in vulnerable groups compared to the resilient groups, but it was counterbalanced between yoked and trained groups. **c** Post- training fear response in Context B (average) was still significantly higher in the vulnerable group. **d** Extinction time curves of resilient subjects no effect of training, although potential floor effect is a confound in the present measurement (left panel). In contrast, training resulted in significantly faster extinction curve in vulnerable subjects (right panel). **e** Volcano plots of differentially expressed genes in the medial prefrontal cortex (mPFC) and hippocampus (HPC) following operant training in vulnerable and resilient groups as compared to yoked controls.

To explore underlying molecular changes underlying this extinction-enhancing effect, we assessed the expression of 45 candidate genes (characterizing basic network functions and involving genes implicated in PTSD risk) in the mPFC and the hippocampus, i.e. two major brain regions altered in PTSD and involved in learning processes (Shin and Handwerger, 2009; Liberzon and Abelson, 2016). Fig.4e and Supplementary Table 1. show the impact of operant training (compared to yoked controls) in the vulnerable and resilient groups. Our exploratory analysis indicated that prefrontal gene expression is more affected compared to the hippocampus, particularly in the resilient group (mPFC: resilient: *Maob* (monoamine oxidase B), *Bcan* (brevican), *Nrxn1* (neurexin 1), *Igf1* (insulin like growth factor 1), *Rtn4r* (Nogo receptor), *Nlgn1* (neuroligin 1), *Fos* (c-fos); vulnerable: *Grm1* (metabotropic glutamate receptor 1); Hippocampus: resilient: *Maoa* (monoamine oxidase A), *Crh* (corticotropin releasing hormone), *Grm1*; vulnerable: *Nlgn1*), and the majority of affected genes is involved in plasticity regulation and learning (Park and Poo, 2013; Kumar et al., 2017; Sudhof, 2018; Thompson et al., 2018).

The potential floor effect observed in the resilient group posed a challenge in determining the extent to which intense plasticity-related changes correlated with network and behavioral functions among resilient individuals. We hypothesize that operant learning recruited mPFC networks that mediated increasing performance *via* plastic changes in the network. Despite the more apparent behavioral outcome (i.e. enhanced extinction), vulnerable subjects exhibited less gene expression alterations, however, the function of these genes is again related to synaptic plasticity (metabotropic glutamate receptor 1 (*Grm1*) and neuroligin-1 (*Nlgn1*)), that can potentially explain the mechanisms of accelerated extinction.

Significantly (uncorrected, p<0.05, blue line) upregulated and downregulated genes (>1.2 fold change, dashed line) are labeled with their names and indicated by red and green, respectively. *Maoa and Maob*: monoamine oxidase A and B; *Bcan*: Brevican; *Igf1*: insulin like growth factor 1; *Nlgn1*: Neuroligin 1; *Ntrk2*: neurotrophic receptor tyrosine kinase 2 (TrkB receptor); *Rtn4r*: Nogo receptor; *Grm1*: metabotropic glutamate receptor 1; *Ngfr*: nerve growth factor receptor (p75NTR). All data are shown as mean ± S.E.M. *p<0.05, ***p<0.001; Two-way ANOVA with Tukey’s post hoc test, or Repeated measures ANOVA.

### Mapping the neuronal activity of the fear network: differential activations in vulnerable and resilient subpopulations

To identify the functional alterations of the fear circuitry, we assessed neuronal activity by means of c-Fos immunolabelling following fear generalization testing (i.e. fear recall in CtxB) (Fig.5a). We focused on specific regions, which have been shown to regulate fear learning and expression (Lopresto et al., 2016; Asok et al., 2018). We found no significant alterations in the activity of the dorsal and ventral subregions of the hippocampus (dorsal: CA1: F_group_(2,23)=1.081, p=0.166; CA3: F_group_(2,23)=0.287, p=0.753; dentate gyrus (DG): F_group_(2,23)=1.233, p=0.31; ventral: CA1: F_group_(2,23)=1.27, p=0.299; CA3: F_group_(2,23)=0.170, p=0.844; DG: F_group_(2,23)=2.753, p=0.084), the paraventricular thalamic nucleus (F_group_(2,20)=1.522, p=0.242), and the basolateral amygdala (F_group_(2,20)=0.845, p=0.444)

**Fig. 5.**
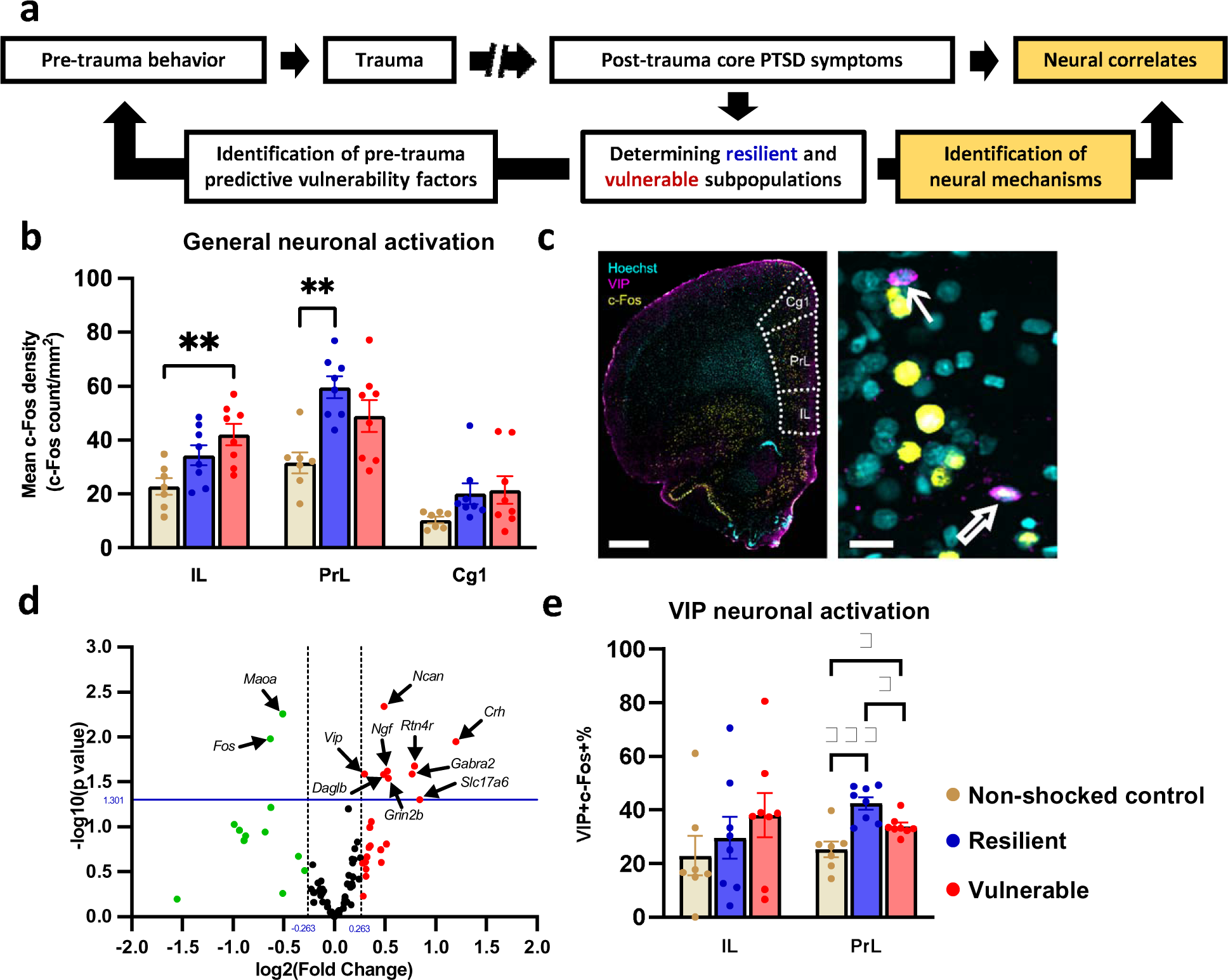
Neural correlates of trauma vulnerability in the mPFC. **a** Experimental design. Data are shown from experimental stages highlighted in yellow. **b** Prefrontal neuronal networks were differentially activated in vulnerable and resilient groups indicated by immunolabeling of c-Fos neuronal activation marker. Whereas infralimbic cortex (IL) showed significant activation in the vulnerable group, prelimbic cortex (PrL) was activated in the resilient group compared to non- shocked subjects. Anterior cingulate cortex (Cg1) activity was not altered by trauma exposure in either group. **c** Representative photomicrographs of Hoechst (blue), Vasoactive intestinal polypeptide (VIP, purple), and c-Fos (yellow) immunohistochemical labeling in the medial prefrontal cortex. Full and empty arrows indicate VIP+ and VIP+/c-Fos+ double labeled neurons, respectively. Scale bars: 1 mm (left), 20 µm (right). **d** Volcano plot of differentially expressed genes. Significantly (uncorrected, p<0.05, blue line) upregulated and downregulated genes (>1.2 fold change, dashed line) are labeled with their names and indicated by red and green, respectively. **e** c-Fos activity of VIP+ neurons in the PrL revealed significantly higher activation in the resilient group. *Crh*: Corticotropin-releasing hormone; *Daglb*: diacylglycerol lipase beta; *Fos*: Fos proto-oncogene, AP-1 transcription factor subunit; *Gabra2*: GABA-A receptor, alpha 2 subunit; *Grin2b*: NMDA receptor 2B subunit; *Maoa*: monoamine oxidase A; *Ncan*: neurocan; *Ngf*: nerve growth factor; *Rtn4r*: Nogo receptor; *Slc17a6*: vesicular glutamate transporter 2; *Vip*: vasoactive intestinal peptide; All data are shown as mean ± S.E.M. *p<0.05, **p<0.01, ***p<0.001, One-way ANOVA with Tukey’s post hoc test.

(Supplementary Fig.3). The central amygdala exhibited significantly reduced activity compared to the resilient group (F_group_(2,20)=4.405, p=0.025; Tukey’s post hoc p=0.024). The mPFC showed significant and differential activation pattern between vulnerable and resilient groups compared to non-shocked controls: whereas the infralimbic cortex (IL) was significantly activated in the vulnerable group only (F_group_(2,20)=6.618, p=0.006; p=0.004 and p=0.082 in vulnerable and resilient groups, respectively), the prelimbic cortex (PrL) exhibited significant activation in the resilient group only (F_group_(2,20)=7.113, p=0.004; p=0.085 and p=0.003 in vulnerable and resilient groups, respectively) (Fig.5b). We did not observe activation in the anterior cingulate cortex (Cg1) (F_group_(2,20)=2.242, p=0.132; Fig.5b). Thus, heightened fear generalization in vulnerable individuals is associated with specific and pronounced changes in neural activation within the fear circuitry, with prominent network operational shifts observed in the mPFC.

### Gene expression correlates of fear generalization in the mPFC

To reveal molecular changes underlying altered mPFC network functioning, we performed an exploratory study and examined expression changes of 92 candidate genes, which have been implicated in PTSD pathogenesis, or characterize basic network functions (Supplementary Table 2). Based on human and preclinical studies of PTSD and fear regulation, we included genes involved in glutamatergic, GABAergic, monoaminergic, cannabinoid, and neuropeptide neurotransmission, as well as neuronal plasticity, HPA-axis function, and inflammatory processes. The volcano plot in Fig.5d shows all differentially expressed genes of the vulnerable group compared to the resilient group. Genes exhibiting significant (p<0.05, uncorrected) changes are indicated by arrows and their abbreviated names. *Fos* (c-Fos: t(13)=2.986, p=0.010) and *Maoa* (monoamine oxydase A: t(13)=3.321, p=0.005) were down- regulated in vulnerable subjects, indicating reduced neuronal activity and lower enzymatic degradation of monoamines. On the other hand, vulnerable subjects exhibited up-regulation of several plasticity-related genes such as *Ncan* (neurocan: t(13)=3.418, p=0.004), *Rtn4r* (Nogo receptor: t(13)=2.621, p<001), *Ngf* (nerve growth factor: (13)=2.556, p=0.023), and genes related to interneuron functions such as *Crh* (corticotropin releasing hormone: t(12)=2.99, p=0.011), *Vip* (vasoactive intestinal peptide: t(13)=2.516, p=0.025), and *Gabra2* (GABA-A receptor, alpha 2 subunit t(13)=2.515, p=0.025), besides other neurotransmission-related genes, i.e. *Grin2b* (NMDA receptor 2B subunit: t(13)=2.461, p=0.028), *Slc17a6* (vesicular glutamate transporter 2: t(13)=2.163, p=0.049), and *Daglb* (diacylglycerol lipase beta: t(13)=2.509, p=0.026). (For exact data set for all genes, see Supplementary Table 2).

### Detailed characterization of mPFC network function: interneuronal activity and monoaminergic inputs

Next, we aimed to assess activity changes in specific interneuron cell types because of their orchestrating function of network activity, and the significant change in *Crh* and *Vip* expression levels that were previously identified as markers of specific interneurons (Klausberger and Somogyi, 2008; Kubota et al., 2011). Corresponding with the latter finding, PrL VIP+ interneurons exhibited significantly higher activity in resilient subjects (F_group_(2,20)=14.15, p<0.001; p=0.032 compared to the vulnerable group) (Fig.5e), which was in concert with higher global PrL activity in resilient subjects as mentioned above (Fig.5b).

Although c-Fos activity of VIP+ interneurons also reflected the directional trend of global c-Fos activity differences across groups in the IL cortex, it was not significantly different between groups (F_group_(2,20)=0.458, p=0.638). In contrast to VIP+ interneurons, no other interneuron types showed significant difference in their activation level between groups in these subregions: parvalbumin-expressing interneurons (PrL: F_group_(2,19)=0.061, p=0.940; IL: F_group_(2,19)=0.684, p=0.516; Supplementary Fig.4a), calretinin-expressing interneurons (PrL: F_group_(2,20)=1.567, p=0.233; IL: F_group_(2,20)=0.243, p=0.786; Supplementary Fig.4b), and somatostatin-expressing interneurons (PrL: F_group_(2,20)=2.860, p=0.080; IL: F_group_(2,20)=2.559, p=0.102; Supplementary Fig.4c). These findings highlight a potential link between mPFC VIP+ interneuron function and altered fear generalization of vulnerable individuals.

Based on decreased *Maoa* gene expression in vulnerable subjects and the first line application of monoaminergic drugs in PTSD and fear-related disorders (Bandelow et al., 2008), we also investigated if vulnerability is associated with altered monoaminergic innervation of the mPFC (Supplementary Fig.5). Using immunostaining of specific synthetizing enzymes, we found no significant alteration of tyrosine-hydroxylase (TH) positive fiber density as an index of dopaminergic inputs (F_group_(2,20)=2.478, p=0.109), dopamine-β-hydroxylase (DBH) positive fiber density as an index of noradrenergic inputs (F_group_(2,20)=1.846, p=0.183), and serotonin transporter (SERT) positive fiber density as an index of serotonergic inputs (F_group_(2,20)=2.499, p=0.107) (Supplementary Fig.5b-d). However, cumulative catecholaminergic density (TH+DBH) showed differential expression, i.e. increased in the resilient group only (F_group_(2,20)=4.537 p=0.023; Tukey’s post hoc: p=0.027 and p=0.079, resilient compared to control and vulnerable groups, respectively), implying variable prefrontal catecholaminergic signaling underlying different outcomes in operant learning and fear generalization.

### Downregulation of prefrontal Crh expression results in prefrontal hyperactivity associated with reduced fear recalls

Elevated expression of prefrontal *Crh* and *Vip*, and the differential activation of VIP+ interneurons in vulnerable subjects implied a potential causal role of interneurons expressing these neuropeptides. Since their significant co-expression has been described in cortical interneurons (Kubota et al., 2011) with further impact of enhanced CRH neurotransmission in anxiety-like behaviour (Toth et al., 2014; Schreiber et al., 2017), we tested their causal role in fear generalization by long-term gene silencing of *Crh* posttrauma using short hairpin RNA (shRNA) (Fig.6a). Injection occurred 2 days after trauma exposure to avoid interference with fear acquisition and memory consolidation. *Crh* shRNA and scrambled RNA (control) groups were counterbalanced for their freezing response exhibited during trauma exposure (U=64, p=0.975). Down-regulation of *Crh* induced significant reduction of fear expression in the traumatic (CtxA: U=2, p<0.001) and safe contexts (CtxB1-B2: U=29, p=0.027; U=16, p=0.001, respectively) (Fig.6b). Latter was associated with marked hyper-activation of mPFC indicated by c-Fos+ cell counts (U=22, p=0.006) (Fig.6c-e). Correlation analysis revealed disrupted prefrontal network function in the *Crh* knocked-down group: whereas the control (scrambled shRNA injected) group showed a positive correlation between prefrontal c-Fos counts and freezing levels (r=0.609, p=0.026) (Fig.6f), this correlation was abolished by *Crh* knock-down (r=0.467, p=0.173) (Fig.6g). Taken together, these findings indicate that disrupted CRH/VIP+ interneuron function has a marked impact on prefrontal network output and related functions such as fear expression in different contexts.

**Fig. 6.**
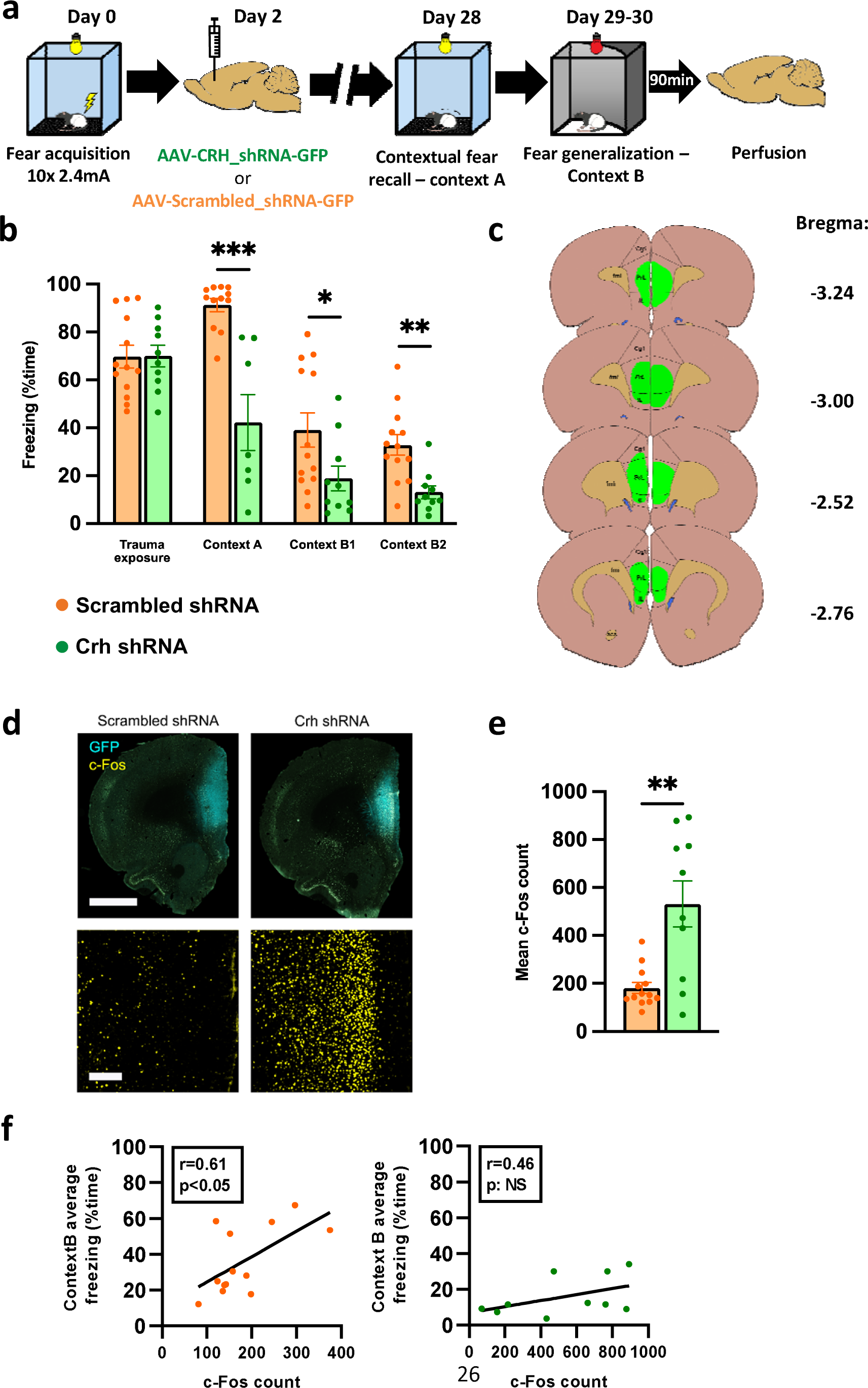
Prefrontal knockdown of corticotropin-releasing hormone (Crh) gene expression reduces long-term fear recall in association with medial prefrontal cortical (mPFC) hyperactivity. **a** Experimental design schematic illustrates bilateral injection of Crh shRNA (short-hairpin RNA) vectors into the medial prefrontal cortex two days after trauma exposure and its impact on long-term fear recalls. Brain samples were taken 90 min after Context B exposure. **b** Crh knockdown significantly reduced long-term fear recall in Context A and B. **c** Coronal brain sections illustrating virus extensions (GFP, green) in the mPFC. **d** Representative photomicrographs illustrating prefrontal c-Fos (yellow) and green fluorescent protein (GFP in cyan) labeling in mPFC from Crh and scrambled shRNA groups. Scale bar: 2 mm (upper), 200 µm (lower). **e** Crh shRNA induced enhanced c-Fos expression in the mPFC compared to scrambled shRNA group. **f** Prefrontal c-Fos counts showed positive correlation with freezing level in Context B in the scrambled shRNA group (left panel), which was not present in the Crh shRNA group (right panel). All data are shown as mean ± S.E.M. **p<0.01, ***p<0.001, Unpaired t-test, Mann-Whitney U test, or Spearman correlation.

## Discussion

The present longitudinal study demonstrates the significant contribution of pretrauma cognitive traits to the development of fear generalization in a rat model of PTSD. Operant learning characteristics predicted individual differences of fear generalization to safe contexts, i.e. vulnerability to developing a core symptom of PTSD. In turn, operant training posttrauma was able to facilitate the extinction of fear generalization in vulnerable subjects suggesting overlapping neural circuits regulating operant and fear learning. Mapping the neuronal activity associated with exaggerated fear generalization (in a safe context) implied the mPFC as a significant mediator: infralimbic and prelimbic cortices exhibited differential activation in vulnerable and resilient groups. This pattern was also reflected in CRH/VIP + interneuron population activity with further alteration of *Crh* and *Vip* mRNA levels in the mPFC. Testing the functional role of these interneurons by means of *Crh* silencing, we observed robust network hyperactivity associated with a reduction of fear expression unspecific to contexts. Taken together, our data suggest that prefrontal network function, reflected in neuronal activity and regulated operant behavior, is a trait-like characteristic that significantly predicts and determines fear generalization following trauma.

First, considering the validity of our model and potential translational interpretations as highlighted by current reviews (Deslauriers et al., 2018; Flandreau and Toth, 2018; Bienvenu et al., 2021), we propose our paradigm as a longitudinal PTSD model designed to dissect specific pretrauma traits and underlying mechanisms that predict and determine fear generalization, a specific core symptom, without covering other symptom domains of PTSD, such as alterations in cognition and mood, and hyperarousal. In this respect, our meta-analysis (Supplementary Fig.1) showed that footshock evokes marked and persistent fear responses related to the traumatic context. The generalization of fear memory exhibited high individual variability that was stable across tests (strong correlation between multiple testing days), suggesting a trait-like characteristic. Importantly, fear generalization was not driven by different fear learning ability (no differences were shown in fear acquisition and contextual recall), hence it provided us a model of maladaptive, progressive and lasting fear generalization. Our model also simulated clinical prevalence (vulnerable quartile) with long-term/persisting outcomes (4 weeks), fulfilling important criteria to model PTSD (Yehuda and Antelman, 1993). An additional advantage of our paradigm was that as a modified version of the Pavlovian fear conditioning paradigm, it is feasible, easily reproducible and can provide interpretable data and comparisons with abundant evidence on short-term conditioned fear mechanisms (Herry and Johansen, 2014).

To date, numerous human studies reported lower cognitive performance in groups diagnosed with PTSD (Macklin et al., 1998; DiGangi et al., 2013), which seem to mediate progression, and determine symptoms trajectories and severity (Kremen et al., 2007; Bardeen et al., 2022; Jagger-Rickels et al., 2022). However, longitudinal studies are still scarce to confirm differences in cognitive performance pretrauma, leaving uncertainty about whether individual variations are predisposing factors or rather consequences of trauma (DiGangi et al., 2013; Jacob et al., 2019). Furthermore, questions persist regarding the particular cognitive domains and associated neurobiological mechanisms that contribute to maladaptive trauma processing. In this respect, our study supports the causal role of lower cognitive function, more specifically, deficits in operant learning and cognitive flexibility as predisposing factors to trauma-induced fear generalization. This cognitive domain was the most powerful in prediction that was replicated in an independent cohort with different settings and task difficulty. In contrast, rigid habitual memory (T-maze performance) as another aspect of cognitive flexibility did not correlate with fear generalization outcome (Schwabe et al., 2010; Goodman et al., 2012). Importantly, this type of habitual inflexibility is distinct from traits associated with set-shifting, including their principal neural elements (striatal-hippocampal vs prefrontal, respectively; (Brown and Tait, 2016)), thereby reinforcing the significance of prefrontal circuits in trauma-induced fear generalization as compared to striatal networks implicated in stress-enhanced habitual memory at the expense of hippocampal-dependent cognitive memory, which may play a role in contextual generalization in PTSD (Goodman et al., 2012). Working memory assessed in the Y-maze, as well as social and contextual-spatial learning abilities were not related to the degree of fear generalization, although their functional contribution to PTSD has been previously proposed (Scott et al., 2015; Liberzon and Abelson, 2016). Our findings overlap with corresponding longitudinal human studies, which reported several cognitive factors as predictive for PTSD outcome, including attention, inhibitory control, flexibility, and executive functions (Samuelson et al., 2020; Schultebraucks et al., 2021), although some domains did not emerge as predictive factors in our model. Outcome measurements may explain these discrepancies since human studies considered PTSD diagnosis as their outcome measure (versions of PCL-5, (Weathers et al., 2018)) in contrast to our animal model focusing on fear generalization specifically.

Longitudinal human studies also showed negative emotionality, anxiety traits, and hyper- arousal as predisposing factors for PTSD (Schultebraucks et al., 2021; Georgescu and Nedelcea, 2023), which was also apparent in our model (Light-Dark Box and Predator odor tests), but with weaker predictive power and effect size compared to cognitive factors. Noteworthy, anxiety tests (Light-Dark Box and Predator odor tests) with more aversive features (high light exposure) showed significant prediction, whereas mildly aversive tests performed under low illumination (Open field and Elevated plus-maze), showed no prediction, suggesting that affective states manifested under elevated stress condition are more related to trauma-related outcomes. We also hypothesize that anxiety and affective traits are stronger determinants of avoidance and hyper- arousal symptom development, which were not assessed in our study as outcomes. In conclusion, we reconcile our data with human finding by hypothesizing separate/specific vulnerability factors or domains (RDoC; (Insel et al., 2010)) as determinants of specific PTSD symptom clusters. Accordingly, intrusive symptoms and fear generalization are significantly determined by operant learning and flexibility related cognitive characteristics.

Latter has implications for clinical interventions, as well. A major clinical goal is to modulate neurocircuits underlying vulnerability using pharmacological and behavioral interventions, or their combination in order to ameliorate symptoms. Latter combinatory approach was a significant endeavour in the last decades exploring pharmacological enhancers of fear extinction-based therapies (Bowers and Ressler, 2015; Singewald et al., 2015). In contrast to pharmacological enhancement, fewer reports are available on the beneficial effect of cognitive trainings as enhancers of fear extinction. These recent studies used cognitive training (e.g. playing Tetris game) either to interfere with memory consolidation of fearful or traumatic stimuli in healthy subjects (Hagenaars et al., 2017), or to blunt intrusive traumatic fear memories in trauma-exposed individuals (Ben-Zion et al., 2018; Iyadurai et al., 2018; Kessler et al., 2018).

Few studies with limited statistical power also showed corrective impact of cognitive training on neurobiological deficits related to PTSD, e.g. hippocampal volume (Butler et al., 2020), supporting our hypothesis that enhanced recruitment of fear-related neurocircuits by cognitive trainings results in plastic changes of these circuits. In line with these findings, our study confirmed posttrauma operant training as a facilitator of fear extinction. Regarding the molecular mediators of this effect, we explored candidate gene expression changes in prefrontal and hippocampal regions and observed altered levels in plasticity-related genes, which have been reported to determine learning abilities and cognitive traits, and have been also implicated in PTSD risk (e.g. metabotropic glutamate receptor 1, (Conquet et al., 1994); neuroligin 1, (Kim et al., 2008; Kilaru et al., 2016)). It supports the hypothesis that cognitive training acts *via* recruitment of neurocircuits which are involved in both operant and extinction learning. By fostering plasticity-related changes, it can lead to modified network operation, which may impede other regulated functions, such as fear extinction.

To explore underlying neurocircuits, we mapped the neuronal activity of several elements of the fear regulatory network relevant to PTSD (Tovote et al., 2015; Ressler et al., 2022). We found the infralimbic and prelimbic regions differentially activated in vulnerable vs. resilient subjects during fear generalization, which corresponds with volumetric and functional deficits of the prefrontal cortex described in PTSD (Shin and Handwerger, 2009; Pitman et al., 2012).

Moreover, the prefrontal cortex is strongly involved in the regulation of both operant learning/cognitive flexibility and fear learning processes (Dejean et al., 2016; Gourley and Taylor, 2016; Howland et al., 2022), positioning it as a prime candidate for mediating the cross- talk between these functions. Accordingly, we further explored the network characteristics of mPFC and found that particularly CRH/VIP+ interneuron population showed alterations, reflected in increased *Crh* and *Vip* mRNA expression in vulnerable subjects that was associated with blunted activity of these neurons (in concert with the whole region) in the prelimbic cortex. We hypothesize that deficient CRH/VIP+ interneuron function resulted in altered network synchronization and regional output in vulnerable subjects based on the significant impact of VIP+ interneurons on network functions (Pi et al., 2013), including during fear expression (Dejean et al., 2016; Karalis et al., 2016). The functional role of VIP+ interneurons across brain regions was shown to integrate relevant sensory stimuli to modulate behavioral adaptation to various challenges, including anxiety and fear learning, mediated by the amygdala, the insular and prefrontal cortices (Krabbe et al., 2019; Lee et al., 2019; Ramos-Prats et al., 2022). Reduced monoaminergic innervation observed in vulnerable subjects suggests that additional factors may contribute to dysregulated prefrontal network function that needs further clarification. Latter is supported by previous studies that highlighted the significant role of monoamines in discrimination performance and cognitive flexibility induced by decreased monoaminergic tones (Clarke et al., 2004; Kehagia et al., 2010).

Finally, our exploratory gene expression analysis revealed changes in both *Vip* and *Crh* expression, pointing out the potential role of this specific cell population in the modulation of fear generalization. Although the neurochemical characteristics of interneuron types are complex, VIP has been shown to co-localize significantly with CRH (Kubota et al., 2011).

Cortical CRH was also shown to modulate anxiety and stress responses in different paradigms (Schreiber et al., 2017), supporting its involvement in fear-related behavior. Previously, enhanced extrahypothalmic/forebrain CRH expression has been shown to increase anxiety, hyperarousal and susceptibility for developing PTSD-like symptoms in a double-hit animal model (Toth et al., 2014; Toth et al., 2016). However, specific targets and mechanisms of these CRH effects are not clarified yet. We aimed to selectively modulate these interneurons to causally test their impact on fear generalization. Although not selectively for fear generalization, we found significant reduction of fear expression after prefrontal *Crh* downregulation. Its association with marked prefrontal hyperactivity can likely traced back to their dysinhibitory function (interneuron-specific interneurons; (Pi et al., 2013)), supporting the central role of CRH/VIP+ interneurons in network synchronization and regional output regulation. The functional coupling between CRH/VIP+ cell activity and fear recall was reflected in their positive correlation in control subjects (scrambled shRNA injected), which was abolished in following *Crh* knockdown resulting in reduced fear expression. This effect suggests that VIP/CRH+ interneurons has marked orchestrating impact on network function, and determines prefrontal output to shape behavioral responses.

Taken together, our unbiased exploration of neurobehavioral vulnerability traits determining trauma-related fear generalization suggests prefrontal cortex dependent cognitive functioning as a significant factor. We observed a strong association between lower pretrauma operant performance and excessive fear generalization, which could be partly reversed by posttrauma operant training leading to facilitated extinction. The degree of fear generalization (vulnerability) was reflected in prefrontal functions on multiple levels: altered gene expression profile related to interneurons (*Crh, Vip*) and plasticity in association with altered network activity. Particularly, CRH/VIP+ interneurons as major coordinators of network activity emerged as an important vulnerability factor for excessive fear generalization. This set of data implies the potential use of prefrontal-dependent cognitive traits as neurobehavioral markers/predictors for fear generalization in PTSD. Such traits may be targeted in preventive measures and cognitive training interventions to alleviate these symptoms.

## Methods

### Animals

Subjects were adult (>10 weeks) male Long-Evans rats (Charles-River Laboratories, Italy), group-housed (4 rats/cage, 60 x 38 x 20 cm) and maintained at controlled temperature of 22±1 °C and relative humidity of 50±10%. Water and laboratory food (Sniff, Germany) were provided *ad libitum*. Subjects were housed on a reverse 12-hour light/dark cycle (on at 8 p.m). All experiments were carried out in accordance with the Directive of the European Parliament and the Council from 22 September 2010 (2010/63/EU) and were reviewed and approved by Hungarian Government Office (PE/EA/874-5/2020), and the Animals Welfare Committee of the Institute of Experimental Medicine.

### Trauma exposure and fear generalization assessment: identification of resilient and vulnerable subpopulations

Subjects were exposed to ten scrambled electric footshocks (2.4 mA; 10 ms pulses interrupted with 20 ms breaks; SuperTech Instruments, Hungary) delivered with 30 s inter-trial intervals (ITIs) during a single 7-min session that started with 2.5 min habituation period. Non-shocked control animals were exposed to the same context without receiving footshocks.

Contextual fear memory was briefly tested 28 days after trauma exposure for 5 min in the same context without delivering footshocks (i.e. CtxA): Plexiglas chamber of 40 x 40 x 40 cm dimensions equipped with a stainless steel electrical grid floor and striped sidewalls (Coulbourn Instruments, Holliston, MA, USA), bright white illumination (300 lux), apparatus cleaned with 20% ethanol between subjects. On the subsequent two days, subjects were exposed to an altered/novel context (CtxB1-2): oval-shaped Plexiglas chambers with plastic floor and solid white sidewalls, red light illumination (5 lux), cleaned with fruity-odor water, conducted by a different experimenter in a different experimental room; i.e. altering all visual, olfactory, and tactile components of the context) for 20 min in order to assess fear expression in a ‘safe’ context (i.e. fear generalization) (Fig.1a-b). The index of fearful response was the time spent with freezing, a species-typical fear response in rodents, quantified with the activity analysis function of Ethovision XT15 software (Noldus, The Netherlands). Experimental populations/cohorts (n=32 or higher) were stratified into quartiles (25%) based on the freezing response exhibited in the two exposures to CtxB to define vulnerable and resilient subpopulations (i.e. high and low fear generalizers). Intermediate quartiles (middle 50%) were not used in our study, except when correlation analysis was conducted on the whole population.

### Behavioral testing

All behavioral test was conducted during the early dark phase, video recorded and analyzed by Ethovision XT15 software. Boxes were cleaned with water and wiped dry between subjects if not stated otherwise.

### Anxiety tests: Light-Dark Box, Elevated Plus-Maze, and Open Field

The Light-Dark Box consisted of a transparent well-lit compartment (50 x 50 x 40 cm, 300 lux) joined to a dark/covered compartment (30 x 50 x 40 cm). Subjects were placed in the light compartment and were allowed to explore for 10 min. The Elevated Plus-Maze was a black arena with 42 x 12 cm arms (35 cm wall height) leveled at 70 cm height and illuminated by dim red light (5 lux). Subjects were placed in the center facing the closed arm and were allowed to explore for 5 min. The Open Field arena was a black non-transparent plastic box (79 x 54 x 35 cm, center defined as 40 x 27 cm) with low intensity illumination (100 lux). Subjects were placed in the corner and were allowed to explore for 10 minutes. In all tests, time spent in the aversive zone, their number of entries and latency to enter were considered as indexes of anxiety. We also combined these variables into composite z-scores for a more reliable index of anxiety.

Total distance moved was the index of locomotor activity.

### Startle response assessment

Startle reactivity was measured in ventilated, sound-attenuating startle chambers (33 x 33x 48 cm; SR-LAB Startle Response System, San Diego Instruments, USA) containing a Plexiglas cylinder (length: 25 cm, d=12 cm). A speaker 24 cm above the cylinder provided background noise (65 dB) and the startle stimuli (20 ms pulses of 80, 90, 100, 110, and 120 dB intensities). The startle session started with a 5 min acclimation period followed by five 120 dB pulses to reach more stable responses. Next, we presented four of each pulse intensities in a pseudo-random order (average 15 s ITIs) to assess average startle response for each intensity.

### Predator odor avoidance test

Innate fear response was assessed by exposing subjects to a synthetic analogue of a fox anogenital odor product, 2-methyl-2-thiazoline (2MT; M83406, Sigma Aldrich) presented in a Plexiglas arena (43 × 27 × 19 cm, 50ul 2MT in a plastic cap at the corner) under bright illumination (300 lux). Subjects were placed in the opposite corner and were allowed to explore for 10 min. Entries and time spent near 2MT source (7 × 11 cm defined zone), and time spent with freezing were indexes of innate fear. We also calculated composite z-score similar to anxiety tests.

### Y-maze

Y-maze was made of three interconnected Plexiglas arms (45 x 17 x 30 cm with 120° angles). Subjects were placed in a dedicated start arm and were allowed to explore for 5 min. Number and sequence of turns made between arms was quantified to calculate the ratio of spontaneous alternation as the index of working memory.

### Social recognition test

The procedure was adapted from Engelmann and his colleagues (Engelmann et al., 2011). Briefly, subjects were habituated in plastic testing cages (60 x 40 x 50 cm, dim illumination) for 2 h before presenting a same-sex juvenile rat (30-35 days old) to interact for 4 min (sampling phase). After 30 min memory retention period, they were introduced to the same familiar juvenile rat and a novel same-sex juvenile rat for 4 min (social recognition phase). Time spent with investigating each juvenile was scored manually by a trained observer. Time preference for the novel juvenile rat was considered as social recognition index.

### Y-place context recognition test

The apparatus was made of transparent Plexiglas arms (each arm: 45 x 17 x 30 cm with 80° angle between choice arms). Choice arms were contextualized by placing distinct objects outside along the walls. Subjects were placed in the start arm and allowed to explore for 10 min with one choice arm closed (sampling phase). After a 30 min memory retention phase spent in their home cage, subjects were re-placed into the start arm and were allowed to explore the arena with two choice arms for 5 min (choice phase). Time preference of novel arm was the index of place recognition.

### Morris Water Maze

The water maze was a black circular pool of 180 cm diameter and 60 cm height filled with 21 °C (room temperature) tap water, which was made opaque by white tempera paint. A white platform of 12 cm diameter was placed in the northwest quadrant and submerged 1.5 cm below the surface. The water maze was surrounded by different visual objects around the pool serving as extra-maze visual cues (under dim illumination, 70 lux). Animals were placed in the pool at one of the five releasing positions for four daily trials for four days. The order of releasing positions was randomized daily but kept constant between subjects. All trials lasted until the animal found the hidden platform (max. 90 s). If the animal did not find the platform during the trial, it was guided to the platform. All subjects were left to stay on the platform for an additional 15 s, then transferred to the next releasing point (start of the next trial). Average daily latency to find the platform (i.e. escape latency) was an index of learning performance. After completing the trials, animals were gently and thoroughly dried by cotton towels before returning to their home cage.

### T-maze

The apparatus was made of three Plexiglas arms (45 x 17 x 30 cm) interconnected in T-shape. Distinct intra-maze cues contextualized the two choice arms: plastic inserts on the walls (dotted vs. striped) and floors (LEGO plate vs. smooth plastic) at the entry zones. The test started with repeated habituation to the maze (5 min daily for 3 days) followed by a training phase (6 trials daily with 30 s it is for 13 days) when one of the choice arms was rewarded by a cornflake (at the end of the arm). The trial ended if the rat turned into either choice arm (and consumed the reward in case of baited arm) or 120 s elapsed without a turn into the arms. Animals were trained to reach the criterion of 5 correct trials (choice of rewarded arm) out of 6. The maze was cleaned with 20% ethanol and wiped dry between trials. In the test phase, the start arm was re-arranged (i.e. moving to the opposite side, from south to north position), while choice arms remained in the same position. Rats were given four test trials to follow either habitual response strategy (turn to the same direction as during training trials) or contextual response strategy (turn based on intra-maze cue, i.e. in opposite direction) in their choices. Frequency of these two strategies was the index of habitual and contextual strategy preference.

### Simple operant learning and Go/No-Go tasks

These tasks were performed in daily training sessions using automated operant chambers equipped with nose poke holes with infrared sensors and LED lights, and a food pellet receptacle in a central position between the nose poke holes (Med Associates, St. Albans, VT, USA). All chambers were located in ventilated sound-attenuating cubicles and controlled by MED-PC IV software. Four days before testing, subjects were put on a restrictive diet (weights maintained at 80-90% of *ad libitum* weight). The task started with two simple operant learning phases (30 min sessions) when LED light cue indicated the rewarded hole. Cue length was 30 s and 10 s during phase 1 and 2, respectively. Rewards were 45mg dustless precision sucrose pellets (BioServ, USA) gained by nose poking into the correct (cued) hole. Correct or incorrect nose pokes terminated the trials, or an omission was registered if no poking occurred. Trials were separated by 5 s intertial intervals (ITIs), when house light turned off to distinguish the active trials from ITIs. Criterion for entering the next phase was >80% accuracy (i.e. the ratio of correct responses) on two consecutive days (applied for all tasks described below). Phase 2 operant task was followed by the Go/No-Go task (40 min sessions), where subjects had to inhibit their response when the No-Go signal (5 s long acoustic cue, 50% of trials in pseudorandom order) was co-presented with the light cue (Go signal for 5 s) to earn reward. A response during No-Go signal (false alarm) terminated the trial without reward. During Go/No-Go task each trial started with a 9-24 s variable pre-cue period when nose pokes during the last 3 s of this period were considered premature responses, which re-started the trial. Registered variables were the total number of responses, percentage of correct and incorrect trials (accuracy), number of premature responses, number of omissions, and days to reach criterion to qualify for the next phase.

### Complex operant learning, strategy set-shifting and 5-choice serial reaction time test

The same operant boxes were equipped with five nose poke holes set on the opposite wall to the food receptacle. Subjects had to learn a more complex operant task (30 min sessions) where the cue light (30 s, illuminating one out of the five nose poke holes in a random manner across trials) was an irrelevant stimulus since the location of the nose poke hole (either 2nd or 4th hole, randomly assigned and counterbalanced within the sample) indicated the rewarded hole. Reward was delivered for correct pokes during cue period or subsequent 10 s limited hold period. House light indicated active trial that was turned off during 5 s ITIs. Operant task was followed by a strategy set-shifting task (30 min sessions) where subjects had to switch from location-based response strategy to a visual cue strategy as the light cue became the relevant stimulus predicting the rewarded nose poke hole position. Finally, subjects entered a 5-choice serial reaction time task (5-CSRTT, based on (Bari et al., 2008)). During this task, subjects had to poke during cue periods (with subsequent 10 s hold-in period) with progressively shortening lengths (from 20 s to 1.25 s) during a 30-min session. Failure to respond during cue or subsequent limited hold periods (omitted trial) or response to non-cued holes (incorrect trial) were registered.

### Posttrauma operant training

One week after fear generalization assessment in CtxB, vulnerable and resilient subpopulations were sorted into ’trained’ and ’yoked control’ groups (counterbalanced for their average freezing level). ‘Trained’ groups were exposed to the same complex operant learning task with five poke holes as described above, whereas ‘yoked controls’ were exposed to operant chambers with random rewarding schedule, i.e. collected the same amount of rewards (median pellet numbers collected by the trained groups on the previous day) that were not dependent on their behavioral performance, but given by a random algorithm. After completing 80% accuracy criterion, subjects were put back on *ad libitum* food access for a week before testing contextual fear recall (5 min in CtxA) and fear generalization (20 min in CtxB) to measure the impact of operant training on fear generalization and extinction.

### Gene expression analysis by quantitative polymerase chain reaction (qRT-PCR)

Immediately, after fear generalization testing in CtxB, rats were decapitated to collect brain samples: 1 mm thick coronal sections were cut (Bregma 3.72 mm to 2.76 mm), and the medial prefrontal cortex was dissected (including the IL, PrL, and Cg1). Samples were snap frozen and kept at -80°C until gene expression analysis. Total RNA was isolated and treated to remove genomic DNA contamination using RNeasy Lipid Tissue Mini Kit (Qiagen, The Netherlands) according to the manufacturer’s protocol. RNA quality and quantity were measured (Agilent 2100 Bioanalyzer, Agilent Technologies, USA or Qubit RNA IQ Assay Kit and Qubit RNA BR Assay Kit, Invitrogen, USA). 1 µg total RNA of each sample was reverse transcribed (High capacity cDNA reverse transcription kit, Thermo Fisher Scientific, USA), and cDNA concentrations were determined (Qubit ssDNA assay kit, Invitrogen, USA). cDNA samples were diluted and combined with TaqMan Gene Expression Master Mix (1 ng/µl final concentration) to create a PCR reaction mix. 92 or 44 candidate genes and four housekeeping genes were analyzed using custom 384-well TaqMan Gene Expression Array Cards (Applied Biosystems, USA).

qRT-PCR was performed (ViiA 7 Real-Time PCR System, Applied Biosystems, USA), and data was collected using the QuantStudio software (Applied Biosystems, USA). Gene expression levels were normalized to the two most stable endogenous control genes across reaction samples (*Actb* and *Gapdh*; geNorm method, Vandesompele, 2002, Genome Biology), and their relative quantity was calculated by the 2^-ddct^ method (Lival, 2001, Methods). Six genes were excluded from the analysis due to low expression levels (Ct numbers >35).

### Immunohistochemical labeling

90 min after fear generalization assessment in CtxB, subjects were deeply anaesthetized (ketamine-xylazine-pipolphen: 200-40-25 mg/kg) and transcardially perfused with cold 0.1 M phosphate-buffered saline (PBS) followed by 4% paraformaldehyde (PFA) in PBS solution. Brains were post-fixed for 3 h, then transferred to 30% sucrose/PBS solution before sectioning 30 µm thick coronal sections using a sliding microtome. Samples were stored at -20 °C in a cryoprotectant solution before immunohistochemical labeling.

First, we labeled c-Fos as an activity marker to map neuronal activity of the fear-regulating network (medial prefrontal cortex, paraventricular thalamic nucleus, central and basolateral nuclei of the amygdala, and CA1-CA3 and dentate gyrus subregions of the dorsal and ventral hippocampus) during fear generalization (CtxB). Second, we co-labeled c-Fos with cell-type specific markers (somatostatin-SST, parvalbumin-PV, calretinin-CR, vasoactive intestinal peptide-VIP), and monoaminergic fibers (serotonin transporter-SERT, tyrosine hydroxylase-TH and dopamine beta hydroxylase-DBH) in mPFC (between Bregma 5.16 mm and 2.52 mm) to characterize network functioning using multiple fluorescent immunolabeling. Briefly, free-floating sections were washed in Tris buffer saline (TBS), and incubated in a blocking solution containing 10% normal donkey serum (NDS; Jackson ImmunoResearch, UK) with 0.3% TritonX-100 in TBS for 1 h. Then slices were incubated in primary antibody solution at 4 °C for 72 h (diluted in TBS with 5% NDS and 0.1% TritonX-100). The following primary antibodies were used: guinea pig anti-c-Fos (1:3000, Synaptic systems, #226-004), mouse anti-NeuN (1:2000, Millipore, #MAB377), rabbit anti-VIP (1:500, Immunostar, #20077), mouse anti-calretinin (1:2000, Swant, #6B3), and rabbit anti-parvalbumin (1:5000, Swant, #PV 27), rabbit anti-somatostatin-14 (1:10000, Peninsula Laboratories International, Inc., #T-4103), anti-TH (1:2000, Millipore, #AB152), mouse anti-DBH (1:2000, Millipore, #MAB308) and guinea pig anti-SERT (1:2000, Synaptic Systems, #340004). Incubation was followed by TBS washes and 2 h long incubation in secondary antibody solution (1:500, with Alexa Fluor 647, Cy3, or Alexa Fluor 488 conjugates, and Hoechst staining in 1:2000 dilution) at room temperature. Finally, slices were washed in TBS, mounted on glass slides, and coverslipped using Mowiol4-88 (Merck).

### Microscopy and image analysis

Slides were imaged using a Pannoramic Digital Slide Scanner (Pannoramic MIDI II; 3DHISTECH, Hungary). An experimenter blind to the experimental groups defined the section planes and the anatomical structures based on reference atlas (Paxinos and Watson, 2007) and manually annotated the mPFC subregions using the CaseViewer 2.4 software (3DHISTECH, Hungary). Total c-Fos signal was counted by using a custom-written ImageJ script. The interneuron marker VIP, CR, PV and SST positive cells and their co-localization with c-Fos signal were manually quantified using the ImageJ software with the Cell counter plugin.

Immunopositive signals (i.e. cell numbers) were counted bilaterally (2-3 sections 360 µm apart) in annotated areas in the above mentioned regions of interest. Signal densities (cell/mm^2^) were calculated and averaged across sections and bilateral areas.

TH, DBH and SERT immunostaining was imaged using a Nikon C2 Confocal Microscope with a 20x objective (Plan Apo VC NA=0.75 WD=1mm FOV=645.12um, Nikon). Three focal planes were captured from the PrL area as z-stack series using the Large area module. Images were converted to maximum-intensity projection images for fiber density analysis, and further processed by background subtraction in ImageJ. Plot Profile function over a standardized area over the cortical layers provided an average value of the intensity of every pixel column (aligned parallel with layers) in our selected area, which was used as an index of fiber density related to every layer.

### Knockdown of Crh expression using shRNA vectors

Adeno-associated virus (AAV) vectors expressing small hairpin RNAs (shRNA) targeting *Crh* or scrambled base sequence (AAV5-EGFP-Scramble_shRNA, #VB010000- 0023jze, scr) as control were purchased from VectorBuilder (Chicago, USA). First, the most effective construct (#VB900052-5912cnc: cnc) was selected from three AAV5-EGFP-rCrh constructs (#VB900052-5919rbk: rbk; #VB900052-5925bpn: bpn; 10e13 GC/ml titers) by injecting each into the mPFC (n=3 animals per group) and quantifying mPFC *Crh* gene expression two and four weeks later by means of TaqMan qRT-PCR using *Crh* primer Rn01462137_m1 (Thermo Fisher), normalized to *Gapdh* expression (Supplementary Fig.6.).

For our behavioral experiment, cnc construct was selected and injected bilaterally into mPFC at the level of PrL and IL cortices (2 x 0.5 µl volume/hemisphere: using AP 2.7 mm, ML 0.5 mm, DV -3.6 and -4.0 mm coordinates from Bregma; Paxinos, 1994) through a glass pipette (tip diameter: 20–30 μm) at a rate of 200 nl/min by using a Nanoject II precision microinjector pump (Drummond, Broomall, PA, USA) under ketamine-xylazine-pipolphen anaesthesia (intraperitoneally, 41,6mg/kg; 8,3mg/kg, 4,16mg/kg, respectively) using stereotaxic equipment (David Kopf Instruments, Tujunga, CA, USA). The pipette was left in place for an additional 5 min after injection to ensure diffusion before slow retraction. After the surgeries, rats received buprenorphine injection (0.1 mg/kg) subcutaneously as analgesic treatment. Surgery was conducted two days after trauma exposure (to avoid interference with acquisition, and to manipulate fear incubation period). Four weeks after virus injection, fear recall tests were conducted as described above. 90 min after fear generalization assessment in CtxB, subjects were anaesthetized and transcardially perfused to verify virus infection sites by means of immunolabeling against the green fluorescent protein (EGFP) imaging (expressed by virus vectors). Only subjects with infection sites limited to mPFC were included in our analysis.

### Statistical analysis

Data are presented as mean ± standard error of the mean. Statistical analysis was performed using GraphPad Prism (GraphPad, USA), Statistica (Tibco, USA), and in R statistical environment. Data were analyzed using Student’s t-test, one- or two-way ANOVA with Tukey’s post hoc test, and repeated measures ANOVA. When the test prerequisites of ANOVA were not fulfilled, we used non-parametric Mann-Whitney U test. Pearson test was used to assess correlations, where data from animals of intermediate quartiles were also included. Distributions and their modality was analyzed by multimode R package (Muller 1991). Statistical significance was set at *p*<0.05 in all cases.

## Acknowledgments

This study was supported by the Hungarian Brain Research Program grant #2017-1.2.1-NKP-2017-00002 (for EM) and the National Research, Development and Innovation Office Grants (Hungary) #K135292 (for EM), #FK129296, #FK142171 (for MT) and #FK128191 (for MA), by the National Laboratory of Translational Neuroscience (#RRF- 2.3.1-21-2022-00011, for EM), and Bolyai Janos Research Fellowship (for MT). We thank all the core facilities of our institute for their supportive help: the Behavioural Studies Unit, Light Microscopy Center, Medical Gene Technology Unit, and Virus Technology Unit. We are grateful to scidraw.io for providing illustrations (doi.org/10.5281/zenodo.3926277; 3926343; 3926015). We also thank Dóra Ömböliné and Brigitta Molnár for their technical assistance.

## Author contributions

L.S. conducted behavioral tests, performed brain sampling, immunohistochemical labeling, microscopy and imaging, data analysis, and wrote manuscript; M.A., G.Y.B., R.D.M., B.Á.V., Z.B., L.B. conducted behavioural tests, performed brain sampling, immunohistochemical labelling, microscopy and imaging with data analysis; Z.K.V. performed statistical analysis, K.D., C.M., H.S. conducted behavioral tests and performed immunohistochemical labelling; Z.B. performed brain surgery and immunohistochemical labelling, imaging with data analysis; G.Y.B., A.S. and A.K. conducted PCR-related sampling and essays, M.T. designed research and experiments, performed brain sampling, immunohistochemical labelling, data analysis, wrote manuscript; É.M. acquired funding, designed research and experiments, performed data analysis, wrote manuscript.

## Competing Interests statement

The authors declare no competing interest.

## Notes

### Competing Interest Statement

The authors have declared no competing interest.

