## Supplementary Material for "Pretrauma cognitive traits predict trauma-induced fear generalization and associated prefrontal functioning in a longitudinal model of posttraumatic stress disorder"

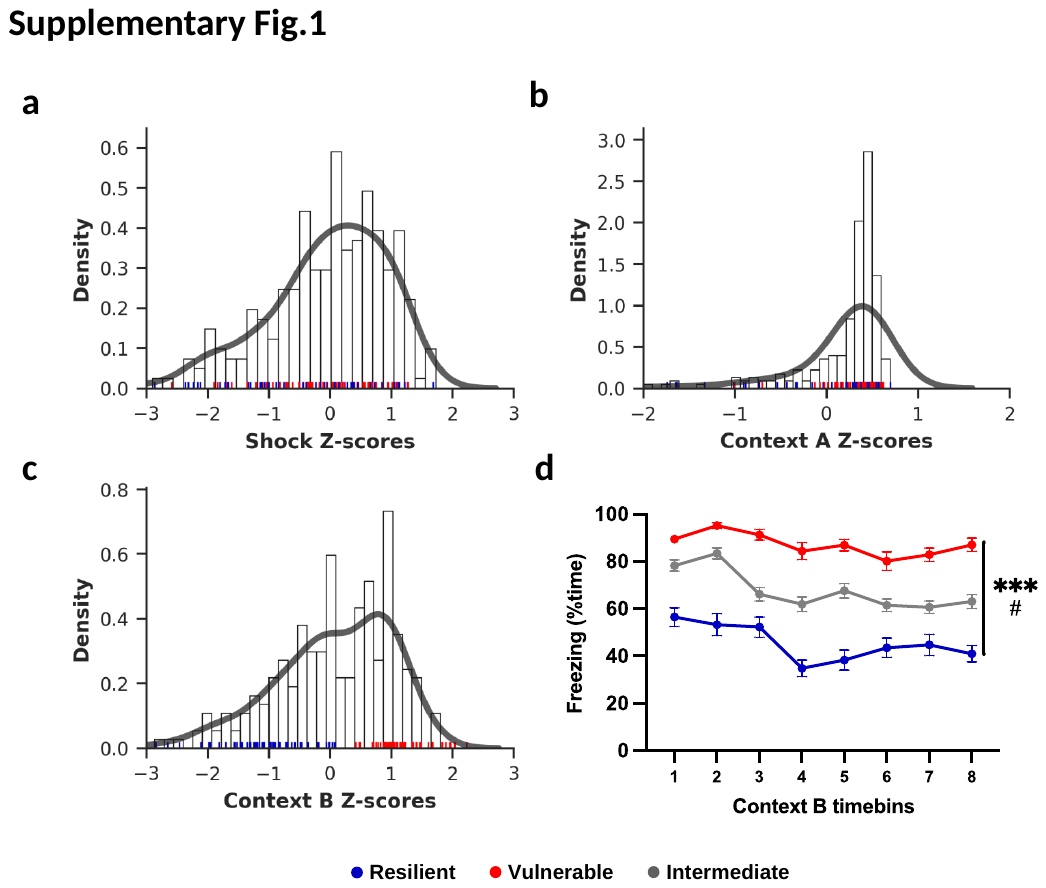

**Supplementary figure 1.** Meta-analysis of standardized freezing levels (z-score of time spent with freezing) from multiple experimental cohorts with the same experimental design (trauma exposure and fear recall testing). Histograms represent freezing z-score distributions of whole populations (including intermediate groups), overlaid by density estimations (thick grey lines). On the x-axes, individual z-scores are presented as rug plots and colored by freezing phenotypes (resilient in blue, vulnerable in red, intermediate subjects are not shown for clarity). a-b Freezing during trauma (footshock) and Context A exposures showed normal distribution with no significant difference between vulnerable and resilient groups as subjects were interspersed along the x axis. **c** Freezing in Context B exhibited clear separation of vulnerable and resilient groups by definition. Moreover, density estimation depicts that distribution is not unimodal, although solid bimodality could not be confirmed. **d** Freezing time curves in Context B points out different lower extinction in the vulnerable groups indicated by significant interaction of time*group besides different freezing levels (generalization). All data presented as mean values ± S.E.M.; p values determined by two-way repeated measures ANOVA with Tukey's post hoc test; ***p<0.001 group effect; #p<0.05 group-time interaction.

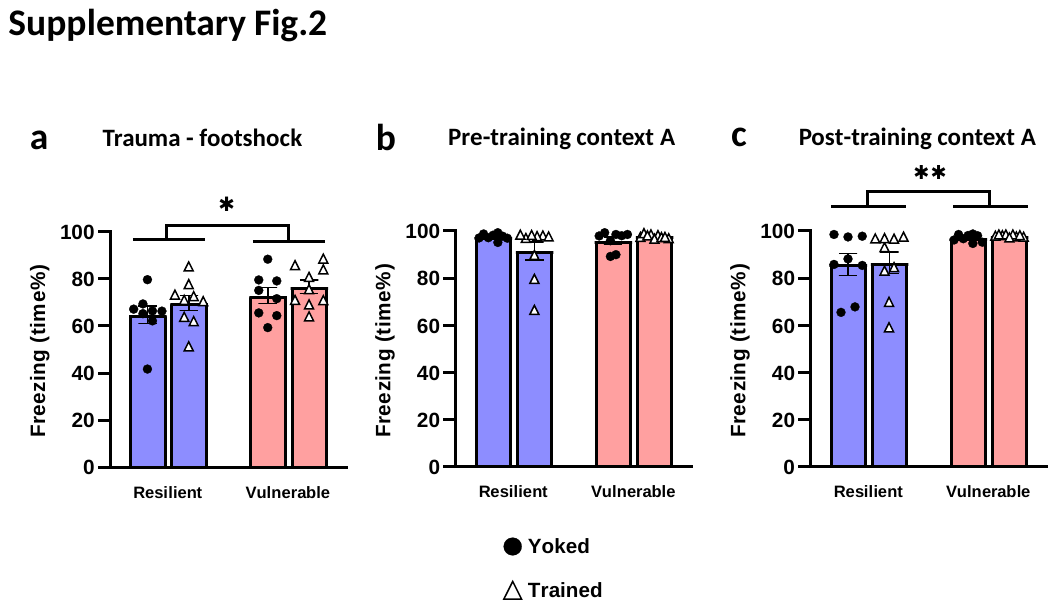

**Supplementary figure 2.** Fear recalls (freezing responses) during trauma exposure, and during exposures to Context A before and after operant training. **a** Vulnerable group showed higher freezing during trauma exposure, although with small effect size. **b** Freezing response was similar between groups during pre-training exposure to Context A, which was significantly higher during post-training exposure to Context A (**c).** Yoked and trained groups exhibited similar fear responses in all tests, i.e. during trauma exposure and both exposures to Context A. All data are shown as mean ± S.E.M. *p<0.05, **p<0.01; Two-way ANOVA with Tukey’s post hoc test.

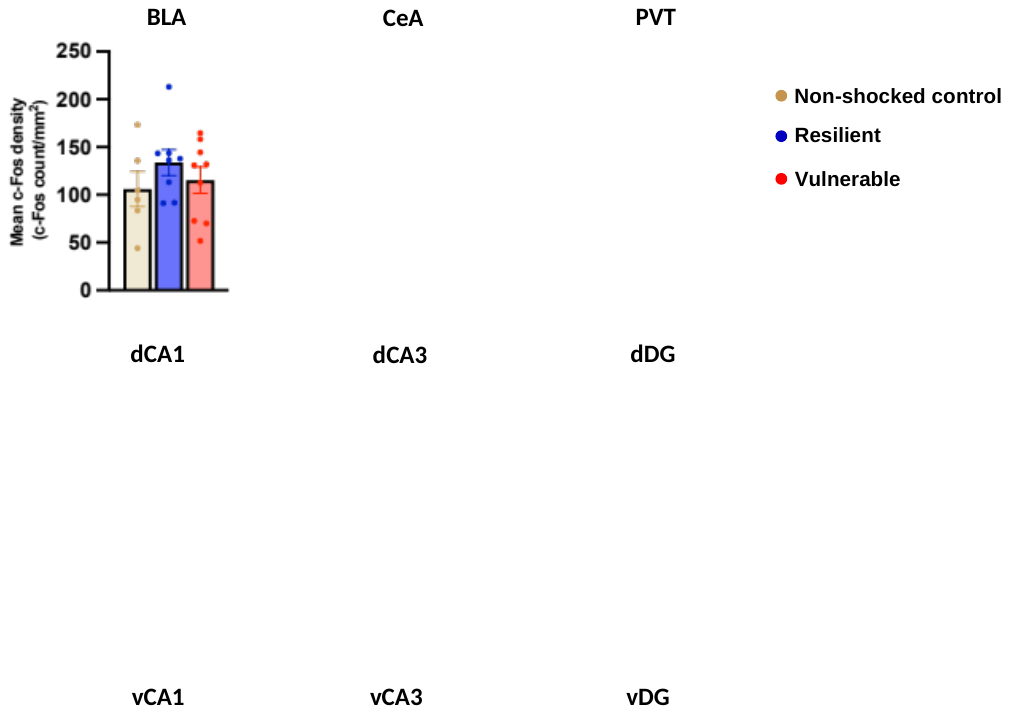

**Supplementary figure 3.** Neuronal activity mapping of the fear circuitry during fear generalization (exposure to Context B). We did not find activity differences between vulnerable and resilient groups in most brain regions except the medial prefrontal cortex (not shown here, see Fig.5), and the central amygdala, where lower activity was observed in vulnerable subjects. BLA: basolateral amygdala; CeA: central amygdala; PVT: paraventricular thalamic nucleus; dCA1-3, vCA1-3: the CA1 and CA3 regions of the dorsal and ventral hippocampus, respectively; dDG, vDG: dentate gyrus of the dorsal and ventral hippocampus, respectively. All data are shown as mean ± S.E.M. *p<0.05, Two-way ANOVA with Tukey’s post hoc test.

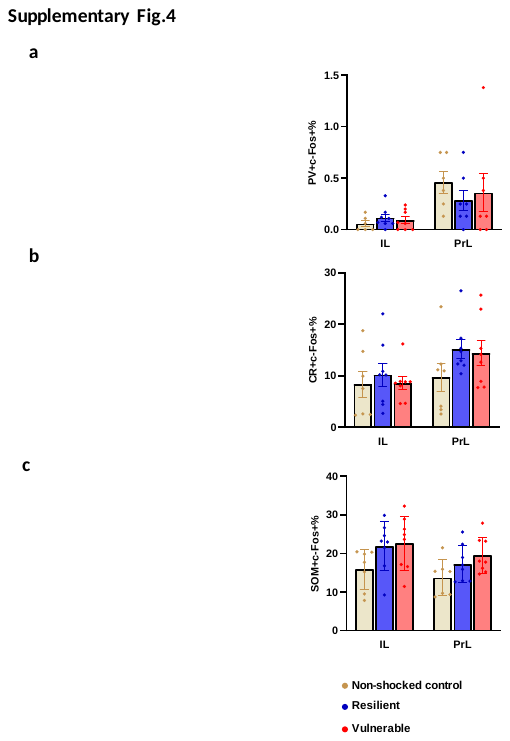

**Supplementary figure 4.** Neuronal activity of different interneuron types in the mPFC during fear generalization (exposure to Context B). Left panels show representative photomicrographs showing immunolabeling of c-Fos with specific interneuron markers (empty arrows), and their co-localization (full arrows). Right panels show quantified of co-localizations, i.e. activity of interneurons positive for **a** parvalbumin (PV), **b** calretinin (CR), or **c** somatostatin (SOM). None of these cell types showed significant difference between groups in the infralimbic (IL), or prelimbic corteces (PrL). Scale bar: 50 µm. All data are shown as mean ± S.E.M. One-way ANOVA.

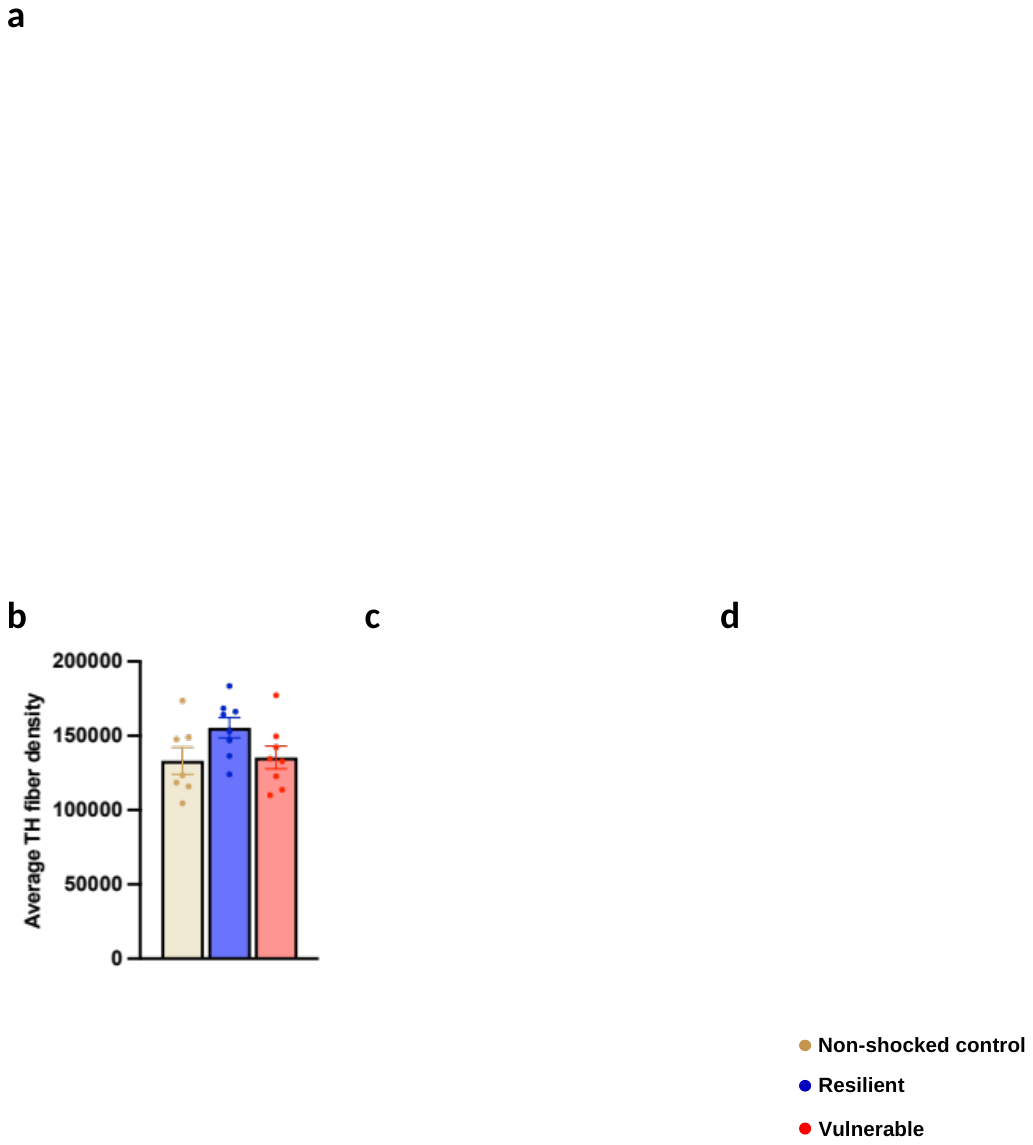

**Supplementary figure 5.** Monoaminergic fiber densities in the medial prefrontal cortex (prelimbic subregion) during fear generalization (exposure to Context B). **a** Representative photomicrographs showing triple labeling of **(b)** tyrosine-hydroxylase (TH), **(c)** dopamine beta-hydroxylase (DBH) and **(d)** serotonin transporter (SERT) as specific markers for dopamine, noradrenaline, and serotonin containing fibers, respectively. Expression density showed no group difference, however, cumulative catecholaminergic density was higher in the resilient group. Scale bar: 500 µm. All data are shown as mean ± S.E.M. One-way ANOVA.

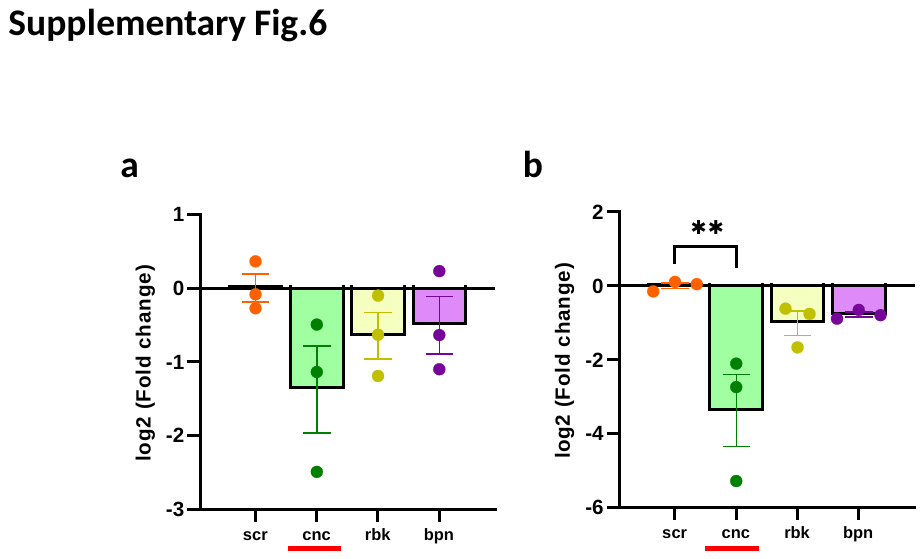

**Supplementary figure 6.** Validation of crh knockdown in the medial prefrontal cortex induced by different AAV-shRNA constructs. Cnc was the most efficient construct to reduce crh expression. Expression was slightly lower at 2 weeks (p=0.081) **(a),** with marked decrease after 4 weeks **(b)**. Other constructs (rbk and bpn) were not effective in the used titers and with this statistical power (n=3). Accordingly, cnc (red underline) with 4 weeks incubation was selected for our behavioral study. All data are shown as mean ± S.E.M. **p<0.01. One-way ANOVA with Tukey’s post hoc test.

**Supplementary Table 1. Gene expression changes in resilient and vulnerable groups following operant training compared to yoked control subgroups.**

| **Medial prefrontal cortex** | | | | | | |
| --- | --- | --- | --- | --- | --- | --- |
| **Resilient** | | |  | **Vulnerable** | | |
| **Gene** | **p value** | **Fold change** |  | **Gene** | **p value** | **Fold change** |
| **Maob** | **0.0001** | **2.367** |  | **Grm1** | **0.016** | **0.279** |
| **Bcan** | **0.0003** | **1.622** |  | Grin1 | 0.111 | 0.746 |
| **Nrxn1** | **0.009** | **4.736** |  | Sst | 0.150 | 1.499 |
| **Igf1** | **0.010** | **2.834** |  | Ngf | 0.176 | 1.389 |
| **Rtn4r** | **0.015** | **1.808** |  | Vip | 0.250 | 0.595 |
| **Nlgn1** | **0.016** | **0.340** |  | Gabra2 | 0.262 | 2.095 |
| **Fos** | **0.018** | **2.037** |  | Nrxn1 | 0.262 | 0.947 |
| Grin2a | 0.057 | 1.832 |  | Bdnf | 0.262 | 0.368 |
| Ncan | 0.069 | 1.527 |  | Nfkb2 | 0.262 | 0.339 |
| Calb1 | 0.081 | 1.395 |  | Drd1 | 0.266 | 0.789 |
| Grin1 | 0.083 | 1.578 |  | Rtn4r | 0.319 | 1.348 |
| Crhr1 | 0.109 | 1.319 |  | Gria2 | 0.327 | 0.720 |
| Gabra2 | 0.148 | 0.709 |  | Fos | 0.331 | 0.739 |
| Maoa | 0.168 | 1.572 |  | Calb1 | 0.367 | 1.233 |
| Fosb | 0.170 | 1.841 |  | Igf1 | 0.423 | 1.026 |
| Pvalb | 0.200 | 0.671 |  | Bcan | 0.457 | 1.156 |
| Acan | 0.221 | 1.711 |  | Grin2b | 0.475 | 0.507 |
| Npas4 | 0.252 | 1.528 |  | Npy | 0.515 | 1.410 |
| Gria1 | 0.262 | 1.302 |  | Daglb | 0.522 | 1.288 |
| Cck | 0.262 | 0.779 |  | Acan | 0.592 | 0.759 |
| Daglb | 0.262 | 0.762 |  | Grin2a | 0.649 | 1.131 |
| Ncam1 | 0.285 | 1.393 |  | Gria1 | 0.667 | 1.095 |
| Gad1 | 0.298 | 1.376 |  | Crhr1 | 0.673 | 1.120 |
| Vip | 0.412 | 1.245 |  | Npas4 | 0.707 | 1.107 |
| Gad2 | 0.423 | 1.220 |  | Maob | 0.743 | 1.104 |
| Gabra1 | 0.423 | 1.061 |  | Arc | 0.745 | 0.908 |
| Grin2b | 0.423 | 0.463 |  | Calb2 | 0.749 | 0.706 |
| Igf2 | 0.489 | 1.256 |  | Gad1 | 0.779 | 1.092 |
| Crh | 0.503 | 0.555 |  | Crh | 0.779 | 0.804 |
| Npy | 0.588 | 0.609 |  | Gad2 | 0.802 | 0.960 |
| Nfkb2 | 0.631 | 2.287 |  | Gabra1 | 0.803 | 0.907 |
| Calb2 | 0.631 | 0.616 |  | Fosb | 0.815 | 1.100 |
| Arc | 0.692 | 1.160 |  | Cck | 0.839 | 1.043 |
| Bdnf | 0.711 | 0.821 |  | Nlgn1 | 0.844 | 1.053 |
| Drd1 | 0.873 | 0.947 |  | Maoa | 0.855 | 0.694 |
| Sst | 0.873 | 0.865 |  | Ncan | 0.861 | 0.958 |
| Ngf | 0.873 | 0.748 |  | Dlg4 | 0.912 | 0.968 |
| Grm1 | 0.924 | 1.058 |  | Igf2 | 0.914 | 0.937 |
| Dlg4 | 1.000 | 0.980 |  | Pvalb | 0.999 | 1.000 |
| Gria2 | 1.000 | 0.601 |  | Ncam1 | 1.000 | 1.216 |
| Ntrk1 | N/A | N/A |  | Ntrk1 | N/A | N/A |
| Ntrk2 | N/A | N/A |  | Ntrk2 | N/A | N/A |
| Oxtr | N/A | N/A |  | Oxtr | N/A | N/A |

| **Hippocampus** | | | | | | |
| --- | --- | --- | --- | --- | --- | --- |
| **Resilient** | | |  | **Vulnerable** | | |
| **Gene** | **p value** | **Fold change** |  | **Gene** | **p value** | **Fold change** |
| **Maoa** | **0.001** | **0.564** |  | **Nlgn1** | **0.006** | **1.226** |
| **Crh** | **0.037** | **0.529** |  | Gad2 | 0.109 | 1.208 |
| **Grm1** | **0.043** | **0.551** |  | Bdnf | 0.150 | 1.362 |
| Grin1 | 0.055 | 0.706 |  | Ngf | 0.200 | 1.178 |
| Ngf | 0.067 | 0.739 |  | Dlg4 | 0.220 | 0.755 |
| Maob | 0.084 | 0.657 |  | Gabra1 | 0.262 | 1.974 |
| Ntrk1 | 0.116 | 0.574 |  | Grm1 | 0.262 | 1.482 |
| Gria2 | 0.145 | 0.707 |  | Npy | 0.262 | 1.406 |
| Nrxn1 | 0.150 | 0.572 |  | Fos | 0.409 | 0.681 |
| Igf2 | 0.206 | 0.583 |  | Fosb | 0.423 | 0.553 |
| Arc | 0.214 | 1.247 |  | Gria2 | 0.429 | 1.202 |
| Ncam1 | 0.288 | 0.760 |  | Ncam1 | 0.479 | 0.848 |
| Cck | 0.324 | 0.819 |  | Ntrk1 | 0.482 | 1.318 |
| Nfkb2 | 0.337 | 0.721 |  | Arc | 0.490 | 0.878 |
| Bcan | 0.360 | 0.818 |  | Vip | 0.508 | 1.188 |
| Pvalb | 0.405 | 1.158 |  | Nfkb2 | 0.508 | 0.853 |
| Gabra1 | 0.423 | 0.826 |  | Bcan | 0.546 | 0.830 |
| Ntrk2 | 0.438 | 0.825 |  | Igf1 | 0.562 | 1.127 |
| Calb2 | 0.462 | 0.885 |  | Acan | 0.562 | 0.862 |
| Npy | 0.465 | 2.085 |  | Calb2 | 0.573 | 1.172 |
| Sst | 0.522 | 1.292 |  | Sst | 0.581 | 1.142 |
| Ncan | 0.525 | 0.849 |  | Crh | 0.613 | 0.887 |
| Daglb | 0.549 | 0.816 |  | Oxtr | 0.631 | 1.176 |
| Drd1 | 0.584 | 1.137 |  | Calb1 | 0.637 | 1.135 |
| Gria1 | 0.593 | 0.871 |  | Grin2a | 0.670 | 0.881 |
| Gabra2 | 0.594 | 0.931 |  | Rtn4r | 0.696 | 0.878 |
| Fosb | 0.631 | 1.364 |  | Gad1 | 0.718 | 0.884 |
| Crhr1 | 0.631 | 1.300 |  | Gabra2 | 0.728 | 0.969 |
| Dlg4 | 0.650 | 0.927 |  | Grin2b | 0.749 | 1.002 |
| Calb1 | 0.670 | 0.917 |  | Maob | 0.753 | 1.101 |
| Gad2 | 0.719 | 1.054 |  | Maoa | 0.781 | 0.895 |
| Grin2b | 0.727 | 0.959 |  | Drd1 | 0.793 | 1.077 |
| Fos | 0.749 | 1.182 |  | Gria1 | 0.795 | 0.934 |
| Igf1 | 0.759 | 0.930 |  | Ncan | 0.819 | 0.932 |
| Npas4 | 0.772 | 1.077 |  | Pvalb | 0.821 | 0.967 |
| Acan | 0.813 | 0.945 |  | Daglb | 0.832 | 0.932 |
| Rtn4r | 0.817 | 0.936 |  | Npas4 | 0.931 | 0.972 |
| Gad1 | 0.844 | 1.055 |  | Ntrk2 | 0.952 | 0.981 |
| Bdnf | 0.873 | 1.165 |  | Nrxn1 | 0.964 | 0.983 |
| Vip | 0.873 | 1.068 |  | Grin1 | 0.968 | 1.012 |
| Oxtr | 0.952 | 1.012 |  | Cck | 0.986 | 0.994 |
| Nlgn1 | 0.969 | 1.007 |  | Crhr1 | 0.988 | 0.995 |
| Grin2a | 0.978 | 0.992 |  | Igf2 | 1.000 | 0.863 |

**Supplementary Table 2. Differentially expressed candidate genes in vulnerable subjects compared to the resilient group.**

| **Gene** | **p** | **Fold Change** |
| --- | --- | --- |
| **Ncan** | **0.005** | **0.241** |
| **Maoa** | **0.006** | **-0.259** |
| **Crh** | **0.011** | **1.440** |
| **Fos** | **0.011** | **-0.398** |
| **Rtn4r** | **0.021** | **0.626** |
| **Ngf** | **0.024** | **0.273** |
| **Gabra2** | **0.026** | **0.589** |
| **Daglb** | **0.026** | **0.239** |
| **Vip** | **0.026** | **0.088** |
| **Grin2b** | **0.029** | **0.284** |
| **Slc17a6** | **0.050** | **0.709** |
| Nfkb2 | 0.061 | -0.392 |
| Maob | 0.063 | 0.019 |
| Npy1r | 0.088 | 0.134 |
| Oxtr | 0.094 | -0.974 |
| Faah | 0.101 | 0.122 |
| Ngfr | 0.109 | -0.878 |
| Bdnf | 0.114 | -0.464 |
| Il1b | 0.125 | -0.765 |
| Adra2b | 0.143 | -0.793 |
| Htr1a | 0.148 | 0.052 |
| Hapln1 | 0.156 | 0.266 |
| Htr3a | 0.162 | 0.128 |
| Grm1 | 0.169 | 0.117 |
| Gria1 | 0.175 | 0.041 |
| Gabra1 | 0.180 | 0.212 |
| Npas4 | 0.213 | -0.126 |
| Adra1d | 0.215 | 0.102 |
| Grm5 | 0.220 | 0.066 |
| Lynx1 | 0.227 | 0.031 |
| Dagla | 0.231 | 0.037 |
| Arc | 0.246 | 0.096 |
| Crhr2 | 0.249 | 0.215 |
| Rtn4 | 0.251 | 0.034 |
| Tnr | 0.254 | 0.078 |
| Cck | 0.264 | -0.045 |
| Adra2a | 0.296 | 0.100 |
| Trpv1 | 0.307 | -0.086 |
| Adra1a | 0.345 | 0.020 |
| Drd5 | 0.354 | 0.097 |
| Htr2a | 0.357 | 0.033 |
| Comt | 0.363 | 0.034 |
| Npy2r | 0.381 | 0.059 |
| Slc17a7 | 0.400 | -0.017 |
| Nr3c1 | 0.420 | -0.026 |
| Cx3cl1 | 0.432 | 0.016 |
| Grin2a | 0.433 | 0.030 |
| Nfkb1 | 0.450 | 0.023 |
| Calb2 | 0.453 | 0.032 |
| Adcyap1r1 | 0.469 | 0.025 |
| Igf1 | 0.490 | -0.019 |
| Acan | 0.493 | -0.052 |
| Ntrk2 | 0.494 | -0.014 |
| Vcan | 0.515 | -0.022 |
| Fkbp4 | 0.516 | -0.011 |
| Nr3c2 | 0.541 | -0.041 |
| Fkbp5 | 0.542 | -0.014 |
| Igf2 | 0.549 | -0.260 |
| Grin1 | 0.581 | -0.018 |
| Il6 | 0.589 | 0.082 |
| Cnr1 | 0.602 | 0.011 |
| Htr2c | 0.636 | -2.413 |
| Dlg4 | 0.646 | 0.011 |
| Adrb2 | 0.684 | 0.016 |
| Pvalb | 0.685 | -0.013 |
| Tnf | 0.689 | -0.041 |
| Mgll | 0.693 | -0.005 |
| Nlgn2 | 0.707 | 0.009 |
| Drd1 | 0.740 | 0.027 |
| Bcan | 0.780 | -0.004 |
| Nlrp3 | 0.806 | 0.008 |
| Nrxn1 | 0.849 | 0.003 |
| Gria2 | 0.853 | 0.002 |
| Ncam1 | 0.868 | 0.002 |
| Adrb1 | 0.882 | -0.001 |
| Slc32a1 | 0.890 | -0.002 |
| Crhr1 | 0.911 | 0.001 |
| Calb1 | 0.918 | -0.001 |
| Lypd1 | 0.929 | 0.000 |
| Htr1b | 0.929 | -0.001 |
| Drd2 | 0.932 | 0.002 |
| Cx3cr1 | 0.936 | -0.001 |
| Nlgn1 | 0.937 | 0.000 |
| Nptx2 | 0.985 | 0.000 |
| Sst | 0.988 | 0.000 |
| Npy | 0.999 | 0.000 |
| Drd4 | N/A | N/A |
| Ifng | N/A | N/A |
| Ntrk1 | N/A | N/A |
| Slc6a2 | N/A | N/A |
| Slc6a3 | N/A | N/A |
| Slc6a4 | N/A | N/A |
